# Disentangling the complex gene interaction networks between rice and the blast fungus identifies a new pathogen effector

**DOI:** 10.1101/2022.07.19.500555

**Authors:** Yu Sugihara, Yoshiko Abe, Hiroki Takagi, Akira Abe, Motoki Shimizu, Kazue Ito, Eiko Kanzaki, Kaori Oikawa, Jiorgos Kourelis, Thorsten Langner, Joe Win, Aleksandra Białas, Daniel Lüdke, Mauricio P. Contreras, Izumi Chuma, Hiromasa Saitoh, Michie Kobayashi, Shuan Zheng, Yukio Tosa, Mark J. Banfield, Sophien Kamoun, Ryohei Terauchi, Koki Fujisaki

## Abstract

Studies focused solely on single organisms can fail to identify the networks underlying host–pathogen gene-for-gene interactions. Here, we integrate genetic analyses of rice (*Oryza sativa*, host) and rice blast fungus (*Magnaporthe oryzae*, pathogen) and uncover a new pathogen recognition specificity of the rice nucleotide-binding domain and leucine-rich repeat protein (NLR) immune receptor Pik, which mediates resistance to *M. oryzae* expressing the avirulence effector gene *AVR-Pik*. Rice Piks-1, encoded by an allele of *Pik-1*, recognizes a previously unidentified effector encoded by the *M. oryzae* avirulence gene *AVR-Mgk1*, which is found on a mini-chromosome. *AVR-Mgk1* has no sequence similarity to known *AVR-Pik* effectors, and is prone to deletion from the mini-chromosome mediated by repeated *Inago2* retrotransposon sequences. AVR-Mgk1 is detected by Piks-1 and by other Pik-1 alleles known to recognize AVR-Pik effectors; recognition is mediated by AVR-Mgk1 binding to the integrated heavy metal-associated (HMA) domain of Piks-1 and other Pik-1 alleles. Our findings highlight how complex gene-for-gene interaction networks can be disentangled by applying forward genetics approaches simultaneously to the host and pathogen. We demonstrate dynamic co-evolution between an NLR integrated domain and multiple families of effector proteins.

## Introduction

Immune recognition between plant hosts and pathogens is often mediated by gene-for-gene interactions [1]. In this classical genetic model, a match between a single plant disease resistance (*R*) gene and a single pathogen avirulence effector (*AVR*) gene leads to pathogen recognition and induces plant immunity [1]. This model is the foundation for understanding *R–AVR* interactions, leading to molecular cloning of numerous *R* and *AVR* genes. However, recent studies revealed there can be a higher level of complexity that expanded the gene-for-gene model [2–5]. In a given plant–pathogen combination, immune recognition frequently involves multiple tangled *R–AVR* interactions. In this case, knockout or knock-in of single host or pathogen genes does not alter the phenotype, hampering attempts to identify genes involved in the interaction. To overcome this problem, we need host and pathogen lines that allow dissection of a single of *R–AVR* interactions. Lines containing only a single *R* or *AVR* locus can be selected from recombinant lines derived from a cross between genetically distant parents. Such materials have been used to analyse the host or pathogen, but have not been simultaneously applied to both the host and pathogen. In this study, we employed integrated genetics approaches on the host and pathogen to unravel complex interactions between rice (*Oryza sativa*) and the rice blast fungus *Magnaporthe oryzae* (syn. *Pyricularia oryzae*).

Studies on the *M. oryzae*–host pathosystem benefited from examining gene-for-gene interactions. The filamentous ascomycete fungus *M. oryzae* causes blast disease in cereal crops, such as rice, wheat (*Triticum aestivum*), and foxtail millet (*Setaria italica*) [6–8]. *M. oryzae* consists of genetic subgroups that have infection specificities for particular host genera [7]. This host specificity is often determined by a repertoire of lineage-specific genes [9–12]. The gain and loss of these lineage-specific genes sometimes results in host jump and specialization [11, 12]. Therefore, identifying host *R* genes with corresponding pathogen *AVR* genes is crucial to understanding host specificities.

Pathogen effectors modulate host cell physiology to promote susceptibility [13]. In *M. oryzae*, at least 15 effector genes have been identified as *AVR* genes [12,14–26]. The protein structures of AVR-Pik, AVR-Pia, AVR1-CO39, AvrPiz-t, AvrPib, and AVR-Pii have been experimentally determined [27–31]. All of their protein structures, except for the zinc-finger fold of AVR-Pii [31], share a similar six-stranded β-sandwich structure called the MAX (*<u>M</u>agnaporthe* <u>A</u>vrs and To<u>x</u>B-like) fold [28, 32]. This sequence-unrelated MAX effector superfamily has expanded in *M. oryzae* and *M. grisea*, probably through diversifying selection and adaptation to the host environment [28,33,34]. Recent advances in protein structure prediction enabled secretome-wide structure prediction to annotate MAX effectors and other effector families in *M. oryzae* [34, 35]. Nonetheless, most MAX effectors remain functionally uncharacterized, including their ability to activate plant immunity.

Similar to other plant pathogenic fungi [36–41], some *M. oryzae* strains contain supernumerary chromosomes called mini-chromosomes (syn. B-, accessory-, or conditionally dispensable chromosomes) in addition to the essential core chromosomes [42–44]. *M. oryzae* mini-chromosomes are smaller than core chromosomes, are rich in transposable elements, and have a lower gene density [45, 46]. *M. oryzae* mini-chromosomes can be hypervariable with frequent inter-chromosomal translocations between core chromosomes and mini-chromosomes [46, 47]. Since mini-chromosomes often carry virulence-related genes, such as *AVR-Pita* [16, 47], *AVR-Pik* [18,46,48,49], a polyketide synthase *Avirulence Conferring Enzyme 1* (*ACE1*) [46, 50], *PWL2* [15, 45], *Biotrophy-associated secreted1* (*BAS1*) [45, 51], and *AvrPib* [23, 45], they are thought to contribute to host adaptation, although the precise mechanisms remain unclear [45–49,52].

To detect invading pathogens, plants evolved disease-resistance genes [53]. Nucleotide-binding domain and leucine-rich repeat protein (NLR) receptors constitute the predominant class of plant intracellular *R* genes [53–55]. The typical domain architecture of plant NLRs is characterized by the central NB-ARC (<u>n</u>ucleotide-<u>b</u>inding adaptor shared by <u>A</u>paf-1, certain *<u>R</u>* genes and <u>C</u>ED-4) domain and the C-terminal leucine-rich repeat (LRR) domain [56]. The N-terminus contains a TIR (<u>T</u>oll/interleukin 1 receptor), CC (Rx-type <u>c</u>oiled-<u>c</u>oil), or CC_R_ (RPW8-type CC) domain [57–59]. NLR genes are often clustered [60], and may consist of a genetically linked pair of NLRs in head-to-head orientation [61–65]. In the prevailing model, NLR pairs consist of functionally specialized sensor and helper NLRs [2,54,65]. Sensor NLRs directly or indirectly recognize pathogen effectors, while helper NLRs are required by sensor NLRs to activate defence signalling. Some sensor NLRs contain non-canonical integrated domains that act as baits for pathogen effectors [66, 67].

In rice, three CC-type NLR pairs, *Pik* (Pik-1/Pik-2), *Pia* (Pia-2/Pia-1, also known as RGA5/RGA4), and *Pii* (Pii-2/Pii-1), have been characterized [61,64,68]. These NLR pairs are genetically linked in head-to-head orientation, and their sensor NLRs (Pik-1, Pia-2, and Pii-2, respectively) have non-canonical integrated domains that mediate pathogen detection. Pik-1 and Pia-2 have a heavy metal-associated (HMA, also known as RATX) domain as the integrated domain [29,64,69]. For Pik-1, the integrated HMA domain, located between the CC and NB-ARC domains, directly binds the *M. oryzae* effectors AVR-PikD, E, and A, and this binding is required to trigger the immune response [29,70–73]. By contrast, the Pia-2 integrated HMA domain C-terminal to the LRR [64] directly binds the two *M. oryzae* effectors AVR-Pia and AVR1-CO39, which have unrelated sequences [69,74,75]. AVR-Pik and AVR-Pik like (APikL) proteins bind members of the host HMA domain family, called small HMA (sHMA) proteins, which may act as susceptibility factors during pathogen infection [33,76–78]. Therefore, the HMA domains of Pik-1 and Pia-2 are considered to act as baits to trap pathogen effectors [66, 67]. Lastly, Pii-2 has an integrated nitrate (NO_3_)-induced (NOI) domain after the LRR domain [79]. Pii-2 indirectly recognizes the *M. oryzae* effector AVR-Pii via a complex between rice EXO70 (a subunit of the exocyst complex) and the NOI domain of Pii-2 [31,79,80]. The integrated domains of these rice sensor NLRs have been used for protein engineering to confer broad-spectrum resistance [81–89].

Since cloning of the NLR pair *Pikm* [61], at least five additional *Pik* alleles (*Pikp*, *Pik*\*, *Pikh*, *Pike*, and *Piks*) have been identified at the *Pik* locus [61,90–95]. This allelic diversification is likely driven by an arms race co-evolution with *M. oryzae AVR-Pik* effectors, where a few Pik amino acid polymorphisms often define their recognition specificity [70–73,96]. The *Pik* alleles, except for *Piks*, were genetically defined as producing resistance against specific isolates of the blast fungus [61,91–95]. However, no report is available for *Piks*-conferred resistance and its target *AVR* gene [96].

In this study, we aimed to uncover additional functions of the well-studied rice Pik immune receptors by integrating host and pathogen genetic analyses (**Fig 1**). This revealed a previously overlooked interaction between a Pik receptor and a *M. oryzae* effector. We found that Piks-1 detects the *M. oryzae* effector AVR-Mgk1, which is unrelated to the AVR-Pik family in sequence and is encoded on a *M. oryzae* mini-chromosome. The integrated HMA domain of Piks-1 binds AVR-Mgk1 but not AVR-PikD, whereas the HMA domains of other Pik-1 alleles bind AVR-PikD and AVR-Mgk1. This study illustrates the potential of integrated host and pathogen genetic analyses to unravel complex gene-for-gene interactions.

**Fig 1.**
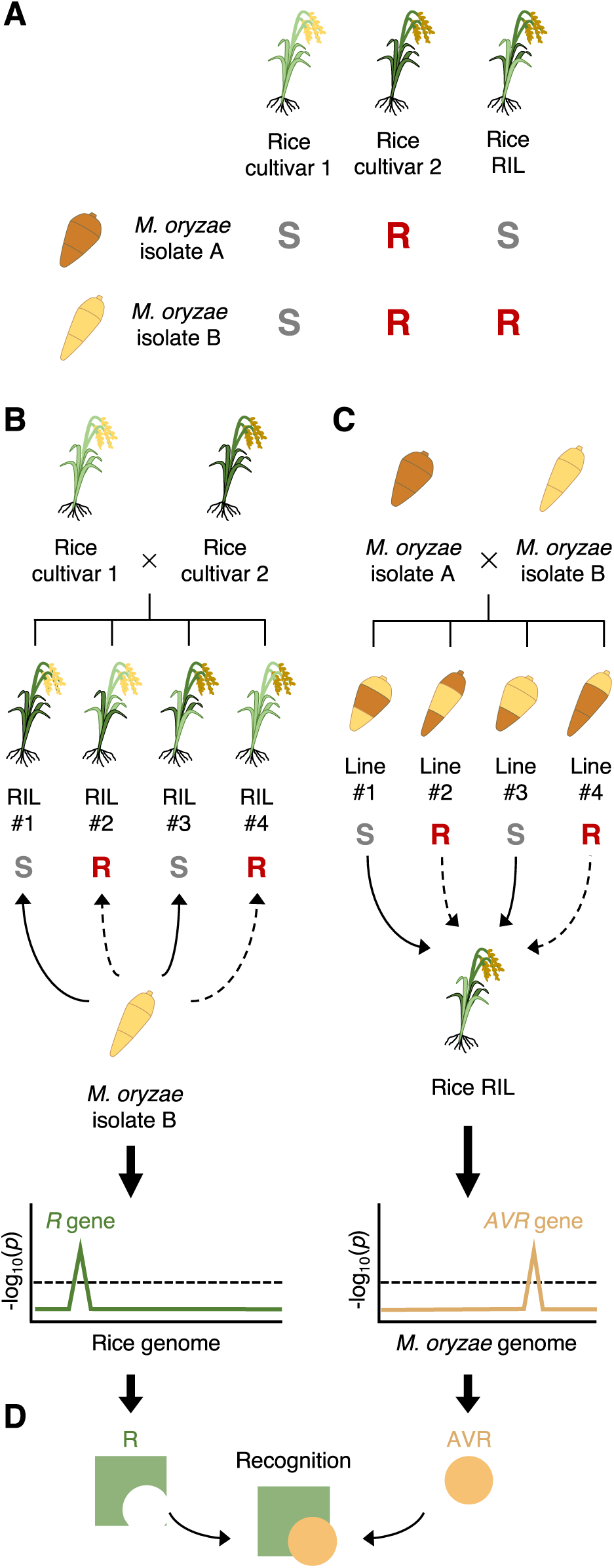
Integrated host and pathogen genetic analyses reveal a previously overlooked gene-for-gene interaction. (A) Rice recombinant inbred lines (RILs) generated to genetically dissect rice resistance to different *M. oryzae* isolates. We generated RILs through self-pollination after the F_1_ generation to reduce heterozygosity. (B) Rice genetics identifies a locus contributing to rice resistance (*R*) to a *M. oryzae* isolate. (C) *Magnaporthe* genetics identifies a locus contributing to avirulence (*AVR*) of a *M. oryzae* isolate to a rice cultivar. (D) Mechanistic studies confirm the gene-for-gene interaction between the identified *R* and *AVR* genes.

## Results

### *Piks* contributes to resistance against *M. oryzae* isolate O23

The *japonica*-type rice cultivar Hitomebore is resistant to the *M. oryzae* isolates TH3o and O23, which originate from Thailand and Indonesia, respectively (**Fig 2A**). In contrast, the *japonica*-type rice cultivar Moukoto is susceptible to these isolates (**Fig 2A**). To determine the loci contributing to the resistance of Hitomebore against TH3o and O23, we produced rice recombinant inbred lines (RILs) derived from a cross between Hitomebore and Moukoto, resulting in 249 RILs that were subsequently subjected to whole-genome sequencing (**S1 Table**). We used 156,503 single nucleotide polymorphism (SNP) markers, designed from the parental genomes, for genetic association analysis on 226 RILs (**S2 Table**). This analysis identified a locus strongly associated with resistance to TH3o on chromosome 1 (**Fig 2B**), and loci associated with resistance to O23 on chromosomes 1 and 11 (**Fig 2C**). The chromosome 1 locus, associated with resistance to both TH3o and O23, contained the NLR gene *Pish*, which confers moderate resistance to *M. oryzae* [97]. In contrast, the locus on chromosome 11 was associated with resistance to O23 only (**Fig 2C**), and contained the NLR gene *Piks*, an allele of *Pik*. A subset of the RILs, including RIL #58, contained the Moukoto-type *Pish* allele and the Hitomebore-type *Piks* allele and was susceptible to TH3o but resistant to O23 (**Fig 2A**), suggesting a role of *Piks* in resistance against O23.

**Fig 2.**
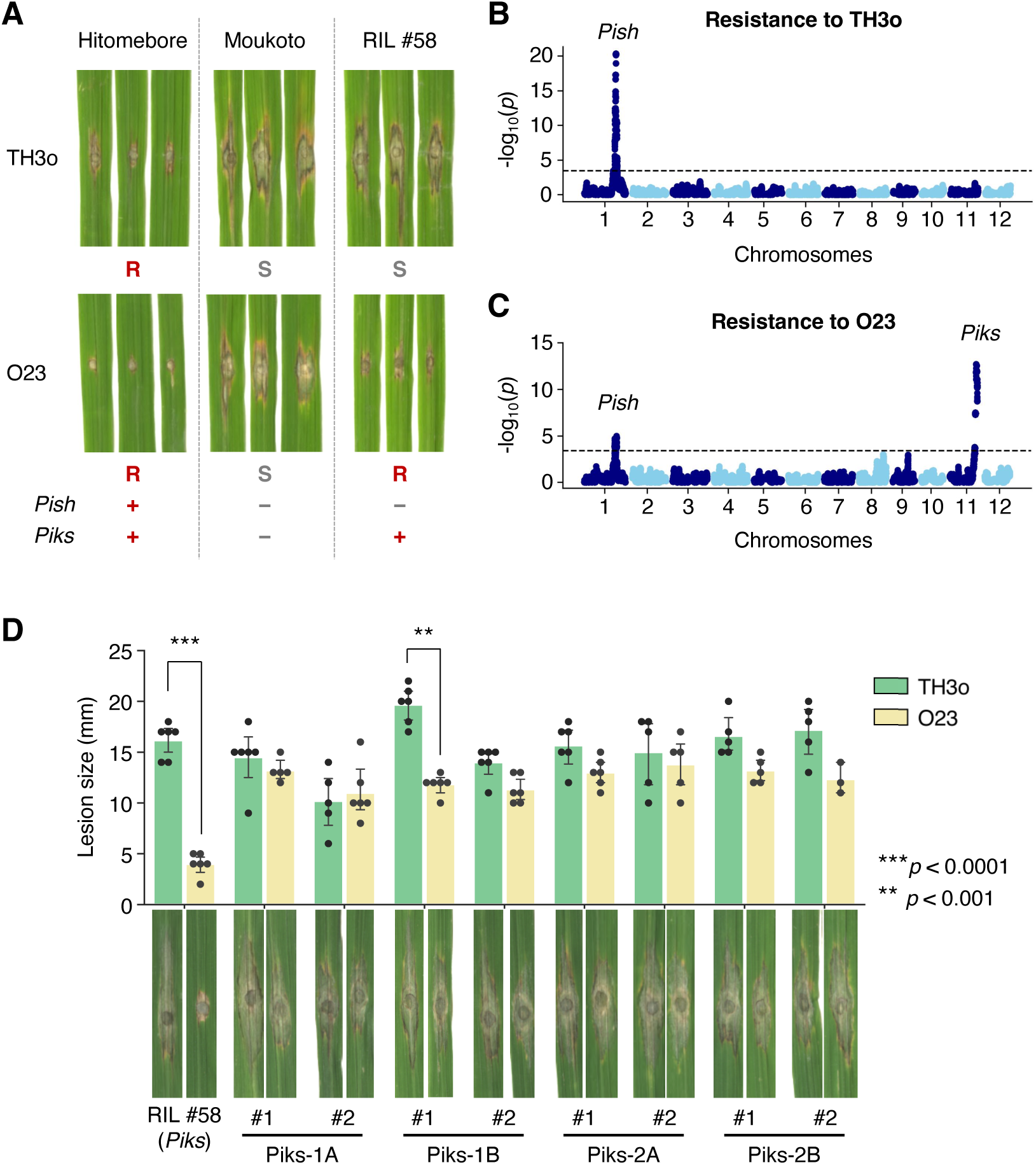
Rice recombinant inbred lines (RILs) untangle the genetics of rice cultivar Hitomebore for resistance to *M. oryzae* isolates TH3o and O23. (A) Punch inoculation assays using *M. oryzae* isolates TH3o and O23 on rice cultivars Hitomebore and Moukoto. Hitomebore is resistant (*R*) to TH3o and O23, while Moukoto is susceptible (*S*) to these isolates. RIL #58, one of the RILs produced from the cross between Hitomebore and Moukoto, is susceptible to TH3o but resistant to O23. (B) Genetic association analysis of rice RIL susceptibility to TH3o identified a locus containing the rice NLR resistance gene *Pish*. (C) Genetic association analysis of rice RIL susceptibility to O23 identified loci containing the rice NLR resistance genes *Pish* and *Piks*. We used 156,503 single nucleotide polymorphism (SNP) markers, designed from the parental genomes, for genetic association analysis on 226 RILs. The vertical axis indicates -log_10_(*p*), where the *p*-value is how likely the marker shows association with a trait due to random chance. The dashed line shows the *p*-value corresponding to a false discovery rate of 0.05. (D) Punch inoculation assays of RNAi-mediated knockdown lines of *Piks-1* and *Piks-2* with the isolates TH3o and O23. We used RIL #58 (*Pish* -, *Piks* +) as the genetic background for the RNAi lines. For each *Pik* gene, we prepared two independent RNAi constructs targeting different regions on the gene (Piks-1A and Piks-1B for *Piks-1*, and Piks-2A and Piks-2B for *Piks-2*, S1 Fig). We performed punch inoculation assays using isolates TH3o and O23 with two RNAi lines per construct, along with RIL #58 as a control. The lesion size was quantified. Asterisks indicate statistically significant differences between TH3o and O23 (two-sided Welch’s t-test). The data underlying Fig 2B–D can be found in S1 Data.

All known *Pik* alleles function as paired NLR genes, consisting of *Pik-1* (sensor NLR) and *Pik-2* (helper NLR), which cooperate to trigger an immune response [61, 98]. Therefore, we performed RNA interference (RNAi)-mediated knockdown of *Piks-1* and *Piks-2* in the RIL #58 (*Pish* -, *Piks* +) background to test their roles in resistance to O23. For both *Piks-1* and *Piks-2*, we targeted two different regions of the open reading frame (**S1 Fig**) and isolated two independent lines per RNAi construct. We used reverse transcription quantitative PCR (RT-qPCR) to analyse *Piks-1* and *Piks-2* expression in these lines (**S2 Fig**). Subsequently, we inoculated the RNAi lines and RIL #58 as a control with the TH3o and O23 isolates (**Fig 2D**). The *Piks-1* and *Piks-2* knockdown lines were susceptible to O23, indicating that *Piks* is involved in resistance to O23.

Although *Pik* is a well-studied NLR gene, the *Piks* allele has not been functionally characterized. Therefore, we investigated the evolutionary relationship of *Piks* and other *Pik* alleles by reconstructing a phylogenetic tree focusing on the *Pik-1* sensor NLRs (**Fig 3A**), which showed that *Piks-1* is most closely related to *Pikm-1*. Comparing amino acid sequences between Piks and Pikm revealed only two amino acid replacements. These two residues were located in the HMA domain of Pik-1 (**Fig 3B**, **S3 Fig**). The HMA domain of Pikm (Pikm-HMA) was crystalized in complex with the *M. oryzae* effector protein AVR-PikD [71]; the two amino acids differentiating Piks-HMA from Pikm-HMA were located at the interface of Pikm-HMA and AVR-PikD (**Fig 3C**), suggesting that these amino acid replacements may affect Pik-1 binding to the AVR-Pik effector. Amino acid sequences of the helper NLRs, Piks-2 and Pikm-2, were identical (**Fig 3B**).

**Fig 3.**
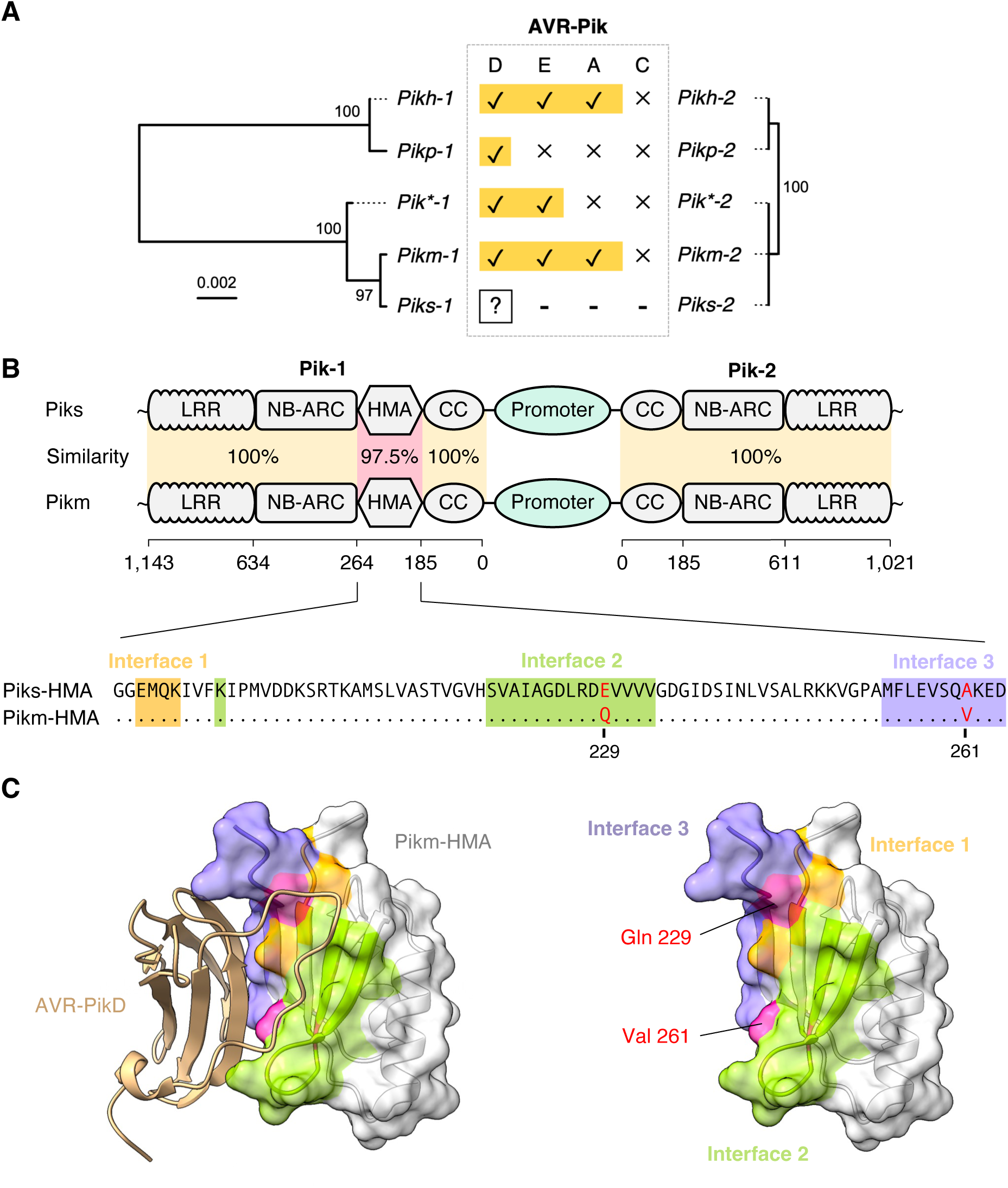
Two amino acid replacements differentiate Piks-1 from Pikm-1. (A) Phylogenetic trees of *Pik* resistance gene alleles are shown together with the experimentally validated protein interactions between Pik and AVR-Pik allelic products. The phylogenetic trees of *Pik-1* and *Pik-2* were drawn based on nucleotide sequences and show the closest genetic relationship between *Piks* and *Pikm*. (B) Schematic representations of the gene locations and domain architectures of the NLR pair genes *Pik-1* and *Pik-2*. The genetically linked Pik-1 and Pik-2 share a common promoter region. Pik-1 has a non-canonical integrated HMA domain that binds *M. oryzae* AVR-Pik allelic products. Piks and Pikm differ by two amino acid replacements located at the integrated HMA domain of Pik-1. These polymorphisms, E229Q and A261V, are located at the binding interface 2 and 3 for AVR-PikD, respectively [71]. We calculated the sequence identities between Piks and Pikm based on amino acid sequences. (C) Structure of Pikm-HMA (PDB ID: 6FU9 chain A) in complex with AVR-PikD (PDB ID: 6FU9 chain B) [71]. The two amino acids differing between Piks-HMA and Pikm-HMA are exposed to the AVR-PikD-interaction site. The colors correspond to the colors of the alignment in (B).

### *Magnaporthe* genetics reveals an avirulence effector gene *AVR-Mgk1* encoded on a mini-chromosome

To identify the *AVR* gene of *M. oryzae* isolate O23 that encodes the effector recognized by Piks, we crossed TH3o and O23 (**Fig 1** and **4A**). We first assembled the genome sequence of O23 into 11 contigs with a total size of 43 Mbp using long sequence reads from Oxford Nanopore Technologies (**S3 Table**). The Benchmarking Universal Single-Copy Orthologs (BUSCOs) value of the assembled genome [99] was 98.2% for the complete BUSCOs using the *Sordariomyceta* odb9 dataset (**S3 Table**). Comparing the O23 assembled contigs with the reference genome version MG8 of *M. oryzae* isolate 70-15 [100] by dot plot analysis revealed that the O23 genome was assembled almost completely end-to-end (**S4 Fig**). Compared to *M. oryzae* isolate 70-15, the O23 genome contained a large rearrangement between chromosome 1 and 6, which has been reported in other *M. oryzae* isolates [45,101–103].

**Fig 4.**
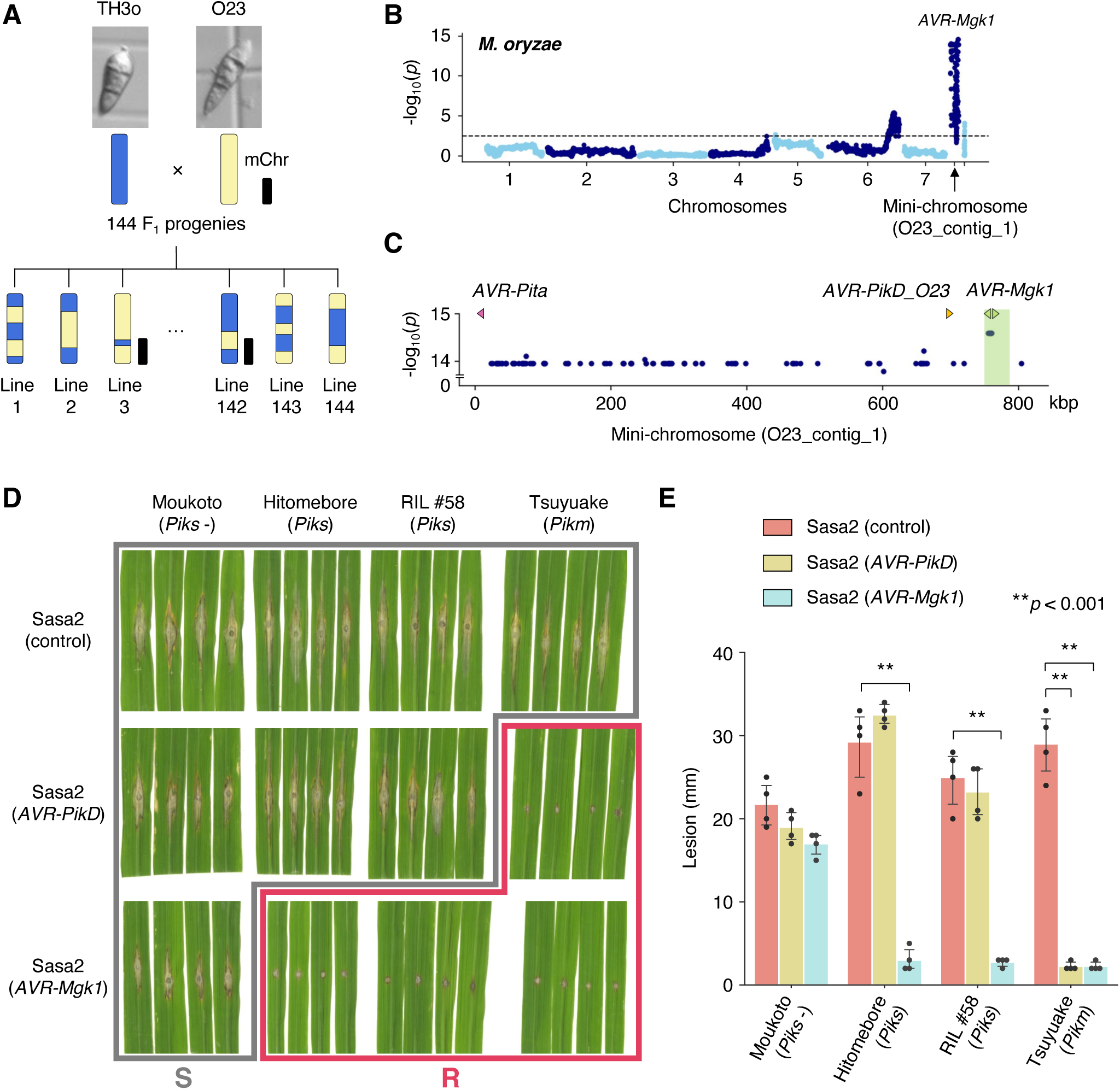
*M. oryzae* genetic analysis identifies an *AVR* gene, *AVR-Mgk1*, encoded on a mini-chromosome. (A) Schematic representations of the F_1_ progeny generated after a cross between *M. oryzae* isolates TH3o and O23. We subjected all F_1_ progeny to whole-genome sequencing. O23 possesses a mini-chromosome [47]. (B) Genetic association of the TH3o x O23 F_1_ progeny using infection lesion size on RIL #58 (*Pish* -, *Piks* +) rice plants as a trait. The vertical axis indicates -log_10_(*p*), where the *p*-value is how likely the marker shows association with a trait due to random chance. The dashed line shows the *p*-value corresponding to a false discovery rate of 0.05. The association analysis based on the O23 reference genome identified *AVR-Mgk1*, encoded on the mini-chromosome sequence O23_contig_1, as an *AVR* gene. O23_contig_1 was not present in the TH3o genome and was unique to the O23 genome. We used 7,867 SNP markers for chromosomes 1–7 and 265 presence/absence markers for the other contigs. (C) *p*-values for O23_contig_1 with annotated *AVR*s. We also detected *AVR-Pita* and *AVR-PikD* in O23_contig_1. *AVR-PikD* in O23_contig_1 contains a frameshift mutation, so we named this variant *AVR-PikD_O23*. The region encoding two *AVR-Mgk1* genes and showing lower *p*-values is highlighted in green. Nucleotide sequences of the two *AVR-Mgk1* genes, arranged in a head-to-head orientation, are identical. (D) Results of punch inoculation assays using *M. oryzae* isolate Sasa2 transformed with *AVR-PikD* or *AVR-Mgk1*. Wild-type Sasa2 infected all the cultivars tested in this study. The Sasa2 transformant expressing *AVR-PikD* infected RIL #58 (*Piks*), but that expressing *AVR-Mgk1* did not infect RIL #58 (*Piks*) or Tsuyuake (*Pikm*) rice plants. (E) Quantification of the lesion size in (D). Asterisks indicate statistically significant differences (*p* < 0.001, two-sided Welch’s t-test). The data underlying **Fig 4B, C, and E** can be found in **S1 Data**.

A study using contour-clamped homogeneous electric field (CHEF) gel electrophoresis identified a mini-chromosome in O23 and reconstructed the sequence of the mini-chromosome region containing the *AVR-Pita* effector [47]. To identify the contigs corresponding to the mini-chromosome in our O23 assembly, we used AVR-Pita as an anchor using the alignment tool Exonerate [104]. AVR-Pita matched the 824-kbp contig named O23_contig_1, which was separately assembled from the core chromosomes (chromosomes 1–7). The presence of the telomeric repetitive sequence TTAGGG [105] in both ends suggested that this contig is a complete mini-chromosome. AVR-Pita was located close to the telomere of the O23_contig_1 as previously reported [47], suggesting that O23_contig_1 likely represents the O23 mini-chromosome [47]. The entire sequence of the O23_contig_1 was absent from the TH3o genome (**S5B Fig**).

We obtained 144 F_1_ progeny from a cross between TH3o and O23 and subjected them to whole-genome sequencing (**S4 Table**). We then compared the TH3o and O23 genome sequences and extracted 7,867 SNP markers for the core chromosomes (chromosomes 1–7) and 265 presence/absence markers for other contigs, including O23_contig_1. Next, we inoculated RIL #58 (*Pish* -, *Piks* +) with each of the *M. oryzae* F_1_ progeny and recorded the lesion size (**S5 Table**, **S6 Fig**). There was a strong association between lesion size and the DNA marker on the mini-chromosome sequence O23_contig_1 (**Fig 4B**). The *p*-values of the DNA markers showing higher levels of association were almost constant across O23_contig_1 (**Fig 4C**), except for position 755–785 kbp with lower *p*-values. This suggested that the candidate *AVR* gene is located on this mini-chromosome region.

To identify the genes expressed within the candidate region, we performed RNA sequencing (RNA-seq) of O23 and TH3o inoculated on barley (*Hordeum vulgare*) cv. Nigrate. Two genes were specifically expressed from the candidate region of O23. These two genes had an identical nucleotide sequence and were arranged in a head-to-head orientation. We named these genes *AVR-Mgk1* (*<u>M</u>a<u>g</u>naporthe* gene recognized by *Pi<u>k</u>*). Sequences similar to *AVR-Mgk1* were not detected in the TH3o genome. These results suggest that AVR-Mgk1 may encode the *M. oryzae* effector recognized by Piks.

To confirm the recognition of AVR-Mgk1 by Piks, we performed a punch inoculation assay using the *M. oryzae* isolate Sasa2, which is compatible with all the cultivars tested in this study, transformed with *AVR-PikD* or *AVR-Mgk1* (**Fig 4D** and **4E**, **S7** and **S8 Figs**). Sasa2 transformants expressing *AVR-PikD* infected RIL #58 (*Piks*) rice plants, but the transformants expressing *AVR-Mgk1* could not (**Fig 4D** and **4E**, **S7 Fig**), indicating that Piks recognizes AVR-Mgk1. Furthermore, Sasa2 transformants expressing *AVR-Mgk1* triggered resistance in the rice cultivar Tsuyuake (*Pikm*). To investigate the recognition specificity of the proteins encoded by other rice *Pik* alleles for AVR-Mgk1, we performed punch inoculation assays with K60 (*Pikp*) and Kanto51 (*Pik\**) rice plants (**S9 Fig**). Sasa2 transformants expressing *AVR-Mgk1* were recognized by K60 (*Pikp*) and Kanto51 (*Pik\**), showing that the proteins encoded by *Pikm*, *Pikp*, and *Pik\** also detect AVR-Mgk1 (**S9 Fig**). These results indicate that AVR-Mgk1 is broadly recognized by Pik proteins.

In addition to *AVR-Mgk1*, we identified a sequence similar to *AVR-PikD* in O23_contig_1 (**Fig 4C**). This *AVR-PikD*-like gene carries a frameshift mutation, and thus encodes a protein with additional amino acids at the C-terminus (**S10A Fig**). We named it *AVR-PikD_O23*. To investigate whether Piks recognizes AVR-PikD_O23, we inoculated RIL #58 (*Piks*) and Tsuyuake (*Pikm*) with Sasa2 transformants expressing *AVR-PikD_O23* (**S10B Fig**). The transformants expressing *AVR-PikD_O23* infected RIL #58 (*Piks*), but not Tsuyuake (*Pikm*) (**S10B Fig**), indicating that AVR-PikD_O23 is not recognized by Piks but is recognized by Pikm, which is consistent with the *AVR* activity of the known *AVR-PikD* gene.

### Retrotransposon repeat sequence-mediated deletion of *AVR-Mgk1* re-establishes virulence

The lower *p*-values of association around the *AVR-Mgk1* genes compared with the rest of the mini-chromosome (**Fig 4C**) facilitated their identification. To identify the F_1_ progeny contributing to the lower *p*-values, we checked the presence/absence of genetic markers on the mini-chromosome in all F_1_ progeny. One F_1_ progeny, named d44a, lacked some markers around the *AVR-Mgk1* genes, suggesting that d44a inherited the mini-chromosome sequence from O23, but lacked the *AVR-Mgk1* genes.

To elucidate the mini-chromosome structure in the d44a isolate, we sequenced the d44a genome using Oxford Nanopore Technologies (**S3 Table**) and compared it with the O23 genome (**Fig 5A**, **S4 Fig**). Two tandemly duplicated sequences of the retrotransposon *Inago2* flanked the *AVR-Mgk1* coding regions in O23. However, in d44a, the *Inago2* sequences were directly associated without the *AVR-Mgk1* coding regions (**Fig 5A**). This suggests that an *Inago2* sequence repeat–mediated deletion of *AVR-Mgk1* occurred in d44a. This deletion was approximately 30 kbp long and the sequence carrying this deletion was assembled separately from the core chromosomes in d44a. This suggests that the deletion was not caused by an inter-chromosome rearrangement between mini- and core chromosomes but by an intra-chromosome rearrangement within or between mini-chromosomes associated with the *Inago2* sequence repeats.

**Fig 5.**
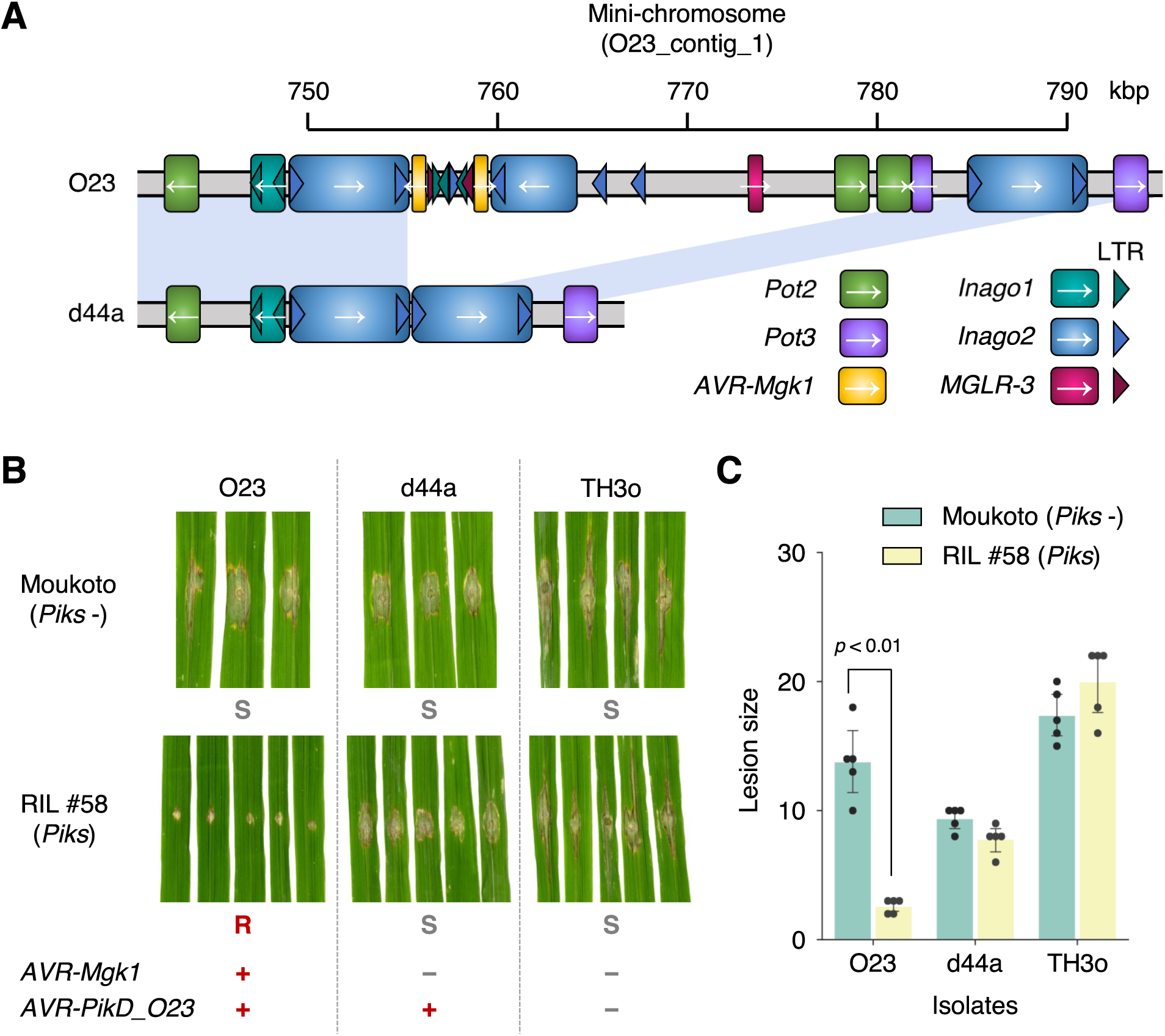
*Inago2* retrotransposon repeat sequence-mediated deletion of *AVR-Mgk1* re-establishes virulence. (A) Comparison of the genomic structures around the *AVR-Mgk1* genes between *M. oryzae* isolates O23 and d44a; d44a is an F_1_ progeny of TH3o x O23. d44a lost the two *AVR-Mgk1* genes. Sequences of transposable elements around *AVR-Mgk1* genes (*Pot2*, *Pot3*, *Inago1*, *Inago2*, and *MGLR-3*) are indicated by color-coded rectangles. Long terminal repeats (LTRs) of retrotransposons are shown in triangles. (B) d44a is virulent against RIL #58 rice plants. We performed punch inoculation assays using O23, TH3o, and d44a on RIL #58 (*Piks*) plants. (C) Quantification of the lesion size in (B). Statistically significant differences are indicated (*p* < 0.01, two-sided Welch’s t-test). The data underlying **Fig 5C** can be found in **S1 Data**.

To investigate the virulence of the d44a isolate on RIL #58 (*Piks*), we performed a punch inoculation assay using O23 and TH3o as controls (**Fig 5B** and **5C**). Consistent with the loss of the two *AVR-Mgk1* genes from the d44a mini-chromosome (**Fig 5A**), d44a infected RIL #58 (*Piks*) plants (**Fig 5B** and **5C**). Since d44a still carries *AVR-PikD_O23* on its mini-chromosome, this result supports that AVR-PikD_O23 is not recognized by Piks.

### AVR-Mgk1 is predicted to be a MAX fold protein that belongs to a distinct family from AVR-Pik effectors

To determine whether AVR-Mgk1 (**Fig 6A**) is related to the AVR-Pik effectors in amino acid sequence, we performed a global alignment between AVR-Mgk1 and AVR-PikD, which revealed a sequence identity of only ∼10% (**S11 Fig**). Therefore, we conclude that these proteins are not related in terms of amino acid sequences.

**Fig 6.**
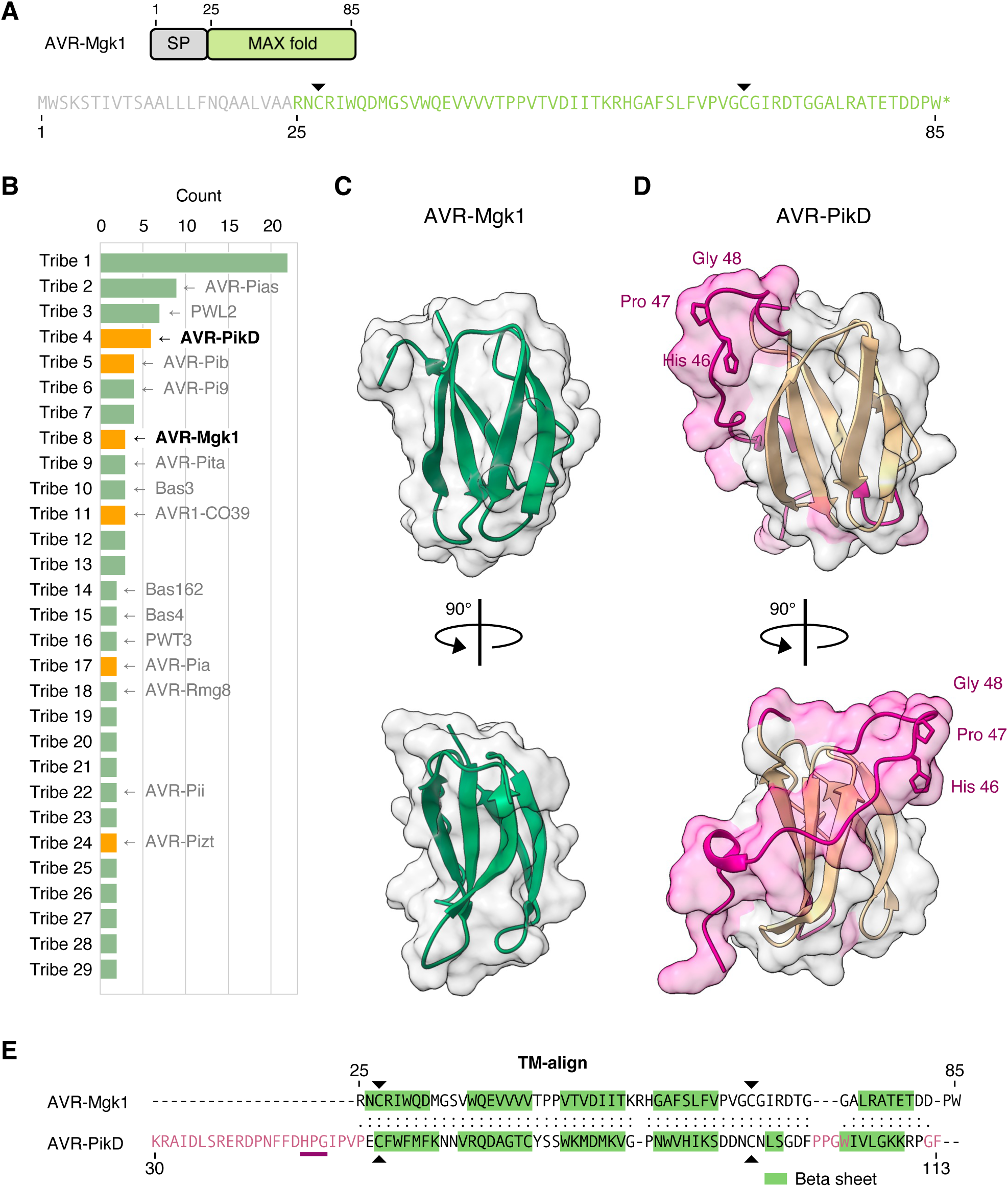
AVR-Mgk1 is predicted to be a MAX fold protein that belongs to a distinct family from AVR-Pik effectors. (A) Domain architecture and amino acid sequence of AVR-Mgk1. We used SignalP v6.0 [106] to predict signal peptide (SP) sequences in AVR-Mgk1. AVR-Mgk1 has the two cysteine residues (Cys27 and Cys67, indicated by black arrowheads) conserved in the MAX effector superfamily. (B) Clustering of putative *M. oryzae* AVR protein sequences using TRIBE-MCL [107]. Tribe-MCL assigned AVR-Mgk1 and AVR-PikD into different tribes. If a tribe includes an experimentally characterized protein, it is shown to represent the tribe. If a tribe includes an experimentally validated MAX effector protein or AVR-Mgk1, the tribe is shown in orange. Tribes having only one protein are not shown. (C) AVR-Mgk1 protein structure predicted by AlphaFold2 [108]. AVR-Mgk1 has antiparallel β sheets, characteristic of the MAX effector superfamily. (D) Protein structure of AVR-PikD (PDB ID: 6FU9 chain B) [71]. (E) Structure-based protein alignment between AVR-Mgk1 and AVR-PikD. TM-align [109] revealed significant structural similarity between AVR-Mgk1 and AVR-PikD, while the regions highlighted in pink structurally differ (C, D). This structural difference involves the highly polymorphic residues (His46-Pro47-Gly48) of AVR-Pik effectors that determine Pik-1 HMA domain binding and are probably modulated by arms race co-evolution [70, 96]. The data underlying **Fig 6B** and **E** can be found in **S1 Data**.

To further investigate the relationship between AVR-Mgk1 and AVR-Pik effectors, we applied TRIBE-MCL clustering algorithm [107] to a dataset of putative *M. oryzae* effector proteins [32], amended with AVR-Mgk1. TRIBE-MCL assigned AVR-Mgk1 and AVR-PikD (**Fig 6B**) into different tribes. This indicates that AVR-Mgk1 belongs to a distinct protein family from AVR-Pik effectors.

Although AVR-Mgk1 has little primary sequence similarity to the AVR-Pik family, AlphaFold2 [108] predicted the protein structure of AVR-Mgk1 as antiparallel β sheets, characteristic of the MAX effector superfamily (**Fig 6C**) [28]. To further evaluate the structural similarity between AVR-Mgk1 and AVR-PikD, we aligned the structures of AVR-Mgk1 (**Fig 6C**) and AVR-PikD (**Fig 6D**) in complex with the HMA domain of Pikm [71] using the structure-based aligner TM-align [109]. TM-align revealed significant structural similarity between the AVR-Mgk1 predicted model and AVR-PikD (**Fig 6E**) with a TM-score >0.5, indicating that they share a similar fold [110]. In addition, AVR-Mgk1 contains the two cysteine residues (Cys27 and Cys67, indicated by black arrowheads, **Fig 6A** and **6E**) conserved in the MAX effector superfamily [28]. Overall, these results indicate that AVR-Mgk1 and AVR-PikD are MAX fold effector proteins that belong to distinct families.

### *AVR-Mgk1* occurs with low frequency in *M. oryzae*

Given that Piks has a narrow recognition spectrum against *M. oryzae* [96], we investigated the distribution of *AVR-Mgk1* in sequenced genomes of the blast fungus. To this end, we performed BLASTN and BLASTP searches against a non-redundant NCBI database using AVR-Mgk1 sequences as query (**S6 Table**). While the BLASTN search failed to find any relevant hits for sequences from the non-redundant nucleotide collection, the BLASTP search found one sequence in the *M. oryzae* isolate Y34 [111] with a sequence identity of ∼52%.

We also performed a BLASTN search against whole-genome shotgun contigs of *Magnaporthe* deposited in NCBI (**S6 Table**). We found sequences identical to *AVR-Mgk1* in the *M. oryzae* isolates 10100 [112] and v86010 [113]. We also found two sequences with ∼91% identity to *AVR-Mgk1* in *M. grisea Digitaria* isolate DS9461 [114], which is a sister species of *M. oryzae* but is genetically markedly different from *M. oryzae* [114, 115]. These results indicate that *AVR-Mgk1* occurs with low frequency in *M. oryzae* and may derive from *M. grisea*.

### The Pik-1 integrated HMA domain binds AVR-Mgk1

The integrated HMA domains of Pia and Pik sensor NLRs (Pia-2 and Pik-1) bind multiple *M. oryzae* MAX effectors [69,75,116]. Therefore, we hypothesized that AVR-Mgk1 binds the integrated HMA domain of Pik-1. To investigate this, we performed yeast two-hybrid assays and *in vitro* co-immunoprecipitation (co-IP) experiments (**Fig 7A** and **7B**). The integrated HMA domain of Pikm-1 bound AVR-Mgk1 and AVR-PikD, whereas the HMA domain of Piks-1 bound only AVR-Mgk1 (**Fig 7A** and **7B**). These results indicate that the Pik-1 integrated HMA domain directly binds AVR-Mgk1, and that one or both of the amino acid changes in Piks-HMA hinder its binding to AVR-PikD (**Fig 3**).

**Fig 7.**
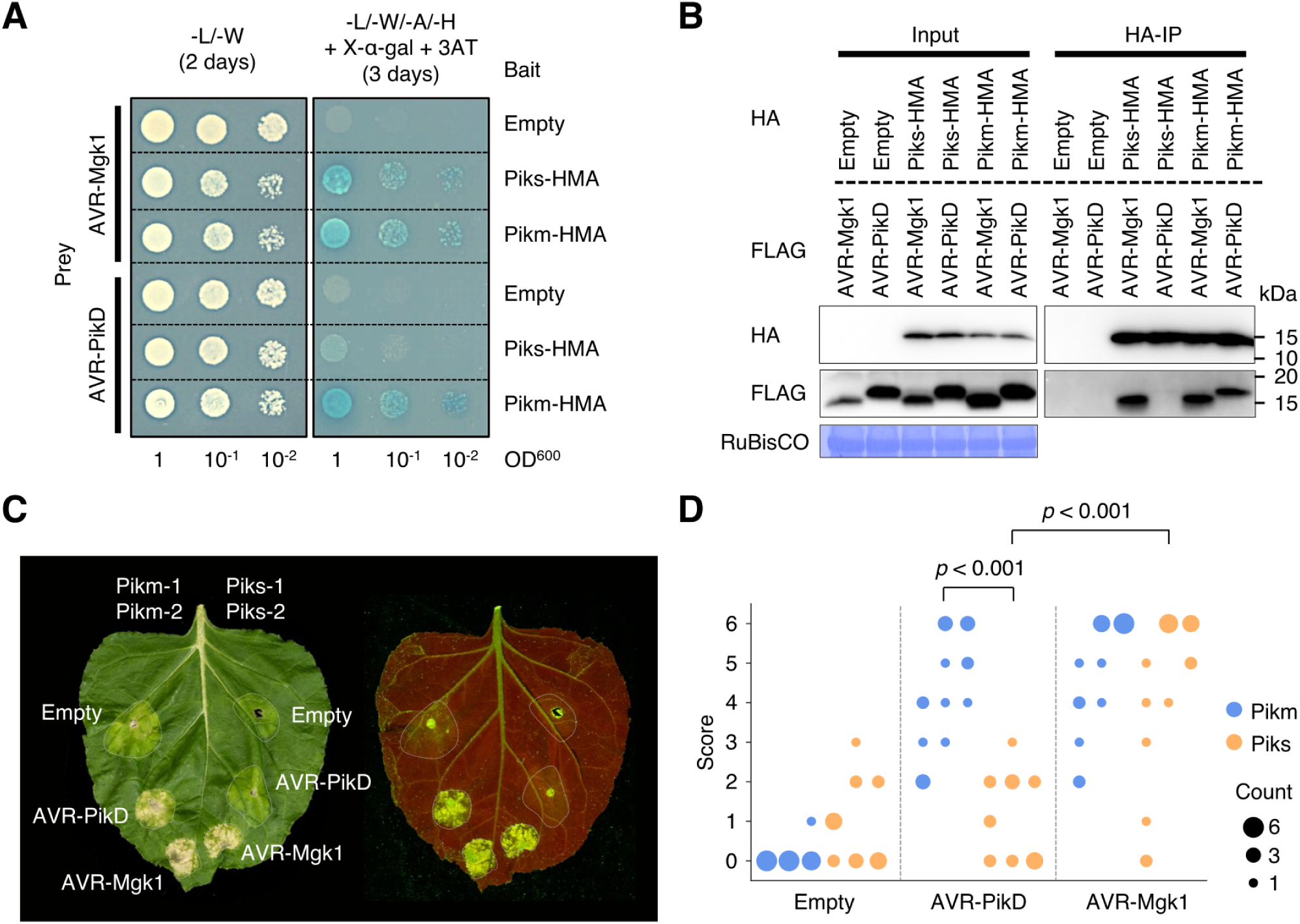
Piks specifically responds to AVR-Mgk1 but not to AVR-PikD. (A) Yeast two-hybrid assays between the Pik integrated HMA domains and AVRs. We used Myc-tagged HMA domains and HA-tagged AVRs as bait and prey, respectively. Empty vector was used as a negative control. Left side: basal medium lacking leucine (L) and tryptophan (W) for growth control. Right side: basal medium lacking leucine (L), tryptophan (W), adenine (A), and histidine (H) and containing X-α-gal and 10 mM 3AT for selection. (B) *In vitro* co-immunoprecipitation (co-IP) experiments between the Pik integrated HMA domains and AVRs. We used N-terminally tagged HA:HMA and FLAG:AVR in the experiments, and the protein complexes were pulled down by HA:HMA using Anti-HA affinity gel. Empty vector was used as a negative control. The large subunit of ribulose bisphosphate carboxylase (RuBisCO) stained by Coomassie brilliant blue is shown as a loading control. (C) Representative images of hypersensitive response (HR) cell death assay after transient co-expression of the AVRs with Pik-1 and Pik-2 in *N. benthamiana*. Pikm and Piks were tested on the left and right sides of the leaf, respectively. The empty vector only expressing p19 was used as a negative control. The leaves were photographed 5–6 days after infiltration under daylight (left) and UV light (right). (D) The HR in (C) was quantified. Statistically significant differences are indicated (Mann-Whitney U rank test). Each column represents an independent experiment. The data underlying **Fig 7D** can be found in **S1 Data**.

To investigate protein-protein interactions between AVR-Mgk1 and the HMA domains of other Pik proteins (Pikp and Pik*), we performed yeast two-hybrid assays and *in vitro* co-IP experiments for Pikp and Pik* (**S12**–**S16 Figs**). The integrated HMA domains of Pikp and Pik* bound AVR-Mgk1 and AVR-PikD, although Pikp bound AVR-Mgk1 with a lower apparent affinity than the other Pik proteins (**S14** and **S16 Figs**). Taken together, these results demonstrated that the HMA domains of all Pik proteins tested bind AVR-Mgk1, which are consistent with the results of the inoculation assay (**S9 Fig**).

### Piks specifically responds to AVR-Mgk1 in a *Nicotiana benthamiana* transient expression assay

The AVR-Pik-elicited hypersensitive response (HR) cell death mediated by Pik NLR pairs has been recapitulated in *Nicotiana benthamiana* transient expression assays [29,71,98]. To investigate whether the HR cell death can be recapitulated with AVR-Mgk1, we performed HR cell death assays in *N. benthamiana* by transiently co-expressing *AVR-Mgk1* or *AVR-PikD* with *Piks* (*Piks-1*/*Piks-2*) or *Pikm* (*Pikm-1*/*Pikm-2*). While Pikm responded to AVR-Mgk1 and AVR-PikD, Piks responded only to AVR-Mgk1 (**Fig 7C** and **7D**). AVR-Mgk1 and AVR-PikD alone did not trigger the HR in *N. benthamiana* (**S17 Fig**). These results are consistent with the protein-protein interaction results (**Fig 7A** and **7B**) and indicate that Piks has a narrower effector recognition range than Pikm.

### Two polymorphisms, E229Q and A261V, between Piks and Pikm quantitatively affect the response to AVR-Pik

We investigated if the amino acid polymorphisms between Piks-1 and Pikm-1 (**Fig 3**) contribute to the differential response to AVR-PikD. We produced single-amino acid mutants of Piks-1 (Piks-1^E229Q^ and Piks-1^A261V^, **Fig 8A**) and performed HR cell death assays in *N. benthamiana* by transiently co-expressing *Piks* (*Piks-1*/*Piks-2*), *Piks^E229Q^* (*Piks-1^E229Q^*/*Piks-2*), *Piks^A261V^* (*Piks-1^A261V^*/*Piks-2*), or *Pikm* (*Pikm-1*/*Pikm-2*) with *AVR-PikD* or *AVR-Mgk1* (**Fig 8B**–**D**). The helper NLRs Piks-2 and Pikm-2 have an identical amino acid sequence (**Fig 3B**). Both polymorphisms (E229Q and A261V) quantitatively affected the response to AVR-PikD (**Fig 8B**). Neither Piks-1^E229Q^ nor Piks-1^A261V^ achieved the same response level as Pikm; however, Piks-1^A261V^ was slightly more responsive to AVR-PikD than Piks-1^E229Q^ (**Fig 8B–D**). The E229Q and A261V mutations did not affect the response to AVR-Mgk1 (**Fig 8C** and **8D**). These results demonstrated that the Q229 and V261 residues of the HMA domain of Pikm are essential for the full response to AVR-PikD.

**Fig 8.**
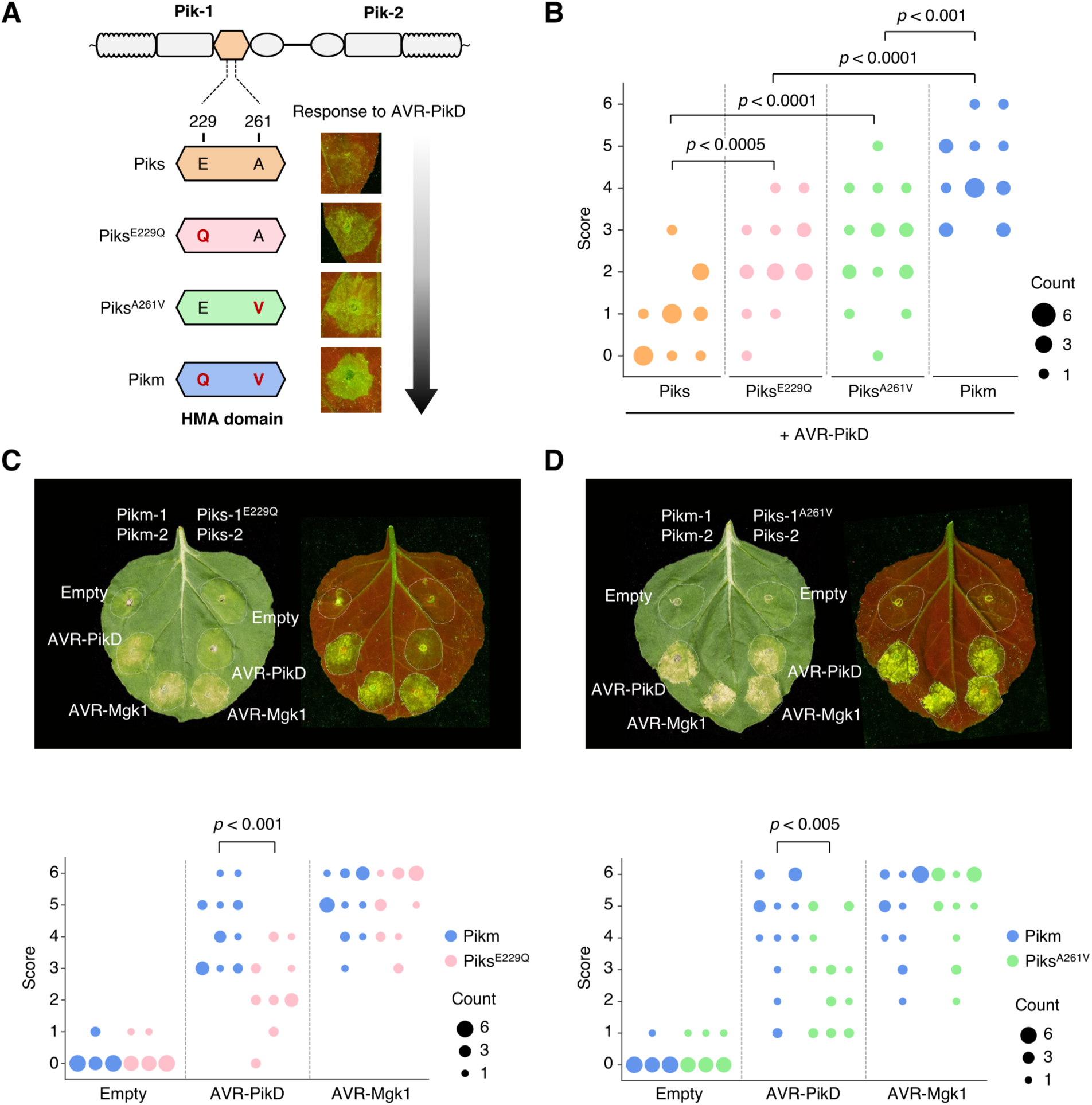
The polymorphisms (E229Q and A261V) between Piks and Pikm quantitatively affect the response to AVR-PikD. (A) Schematic representations of single amino acid mutants (Piks^E229Q^ and Piks^A261V^) used in the HR cell death assay in *N. benthamiana* with AVR-PikD. (B) We quantified HR scores of Piks (Piks-1/Piks-2), Piks^E229Q^ (Piks-1^E229Q^/Piks-2), Piks^A261V^ (Piks-1^A261V^/Piks-2), or Pikm (Pikm-1/Pikm-2) with AVR-PikD 5–6 days after agroinfiltration and statistically significant differences are indicated (Mann-Whitney U rank test). Piks-2 and Pikm-2 are identical. (C) HR cell death assay with Piks^E229Q^ and AVRs. (D) HR cell death assay with Piks^A261V^ and AVRs. The leaves were photographed 5–6 days after infiltration under daylight (left) and UV light (right). We quantified the HR at 5–6 days after agroinfiltration and statistically significant differences are indicated (Mann-Whitney U rank test). Each column represents an independent experiment. The data underlying **Fig 8B**–**D** can be found in **S1 Data**.

To confirm the effects of E229Q and A261V mutations in the HMA domain of Piks-1 on the interaction with AVR-PikD and AVR-Mgk1, we performed yeast two-hybrid assays. The yeast two-hybrid assay showed that both Piks^E229Q^-HMA and Piks^A261V^-HMA as bait bound AVR-PikD as prey to similar levels compared to Pikm-HMA binding with AVR-PikD (**Fig 9A**, **S18 Fig**). This result is consistent with the result of yeast two-hybrid assay in a recent study [117]. On the other hand, we found that Piks^E229Q^-HMA and Piks^A261V^-HMA as prey weakly bound AVR-PikD as bait, compared to Pikm-HMA binding with AVR-PikD (**Fig 9B**, **S19 Fig**). The E229Q and A261V mutations in the HMA domain of Piks-1 did not affect the binding to AVR-Mgk1 (**Fig 9**, **S18** and **S19 Figs**). These results support our findings in HR cell death assay showing that both E229Q and A261V quantitatively affect the response to AVR-PikD but not to AVR-Mgk1 (**Fig 8**). A recent study independently showed the quantitative binding affinity of Piks-HMA mutants (Pikm-HMA > Piks^A261V^-HMA > Piks^E229Q^-HMA > Piks-HMA) to AVR-PikD by analytical gel filtration [117]. Overall, both Q229 and V261 residues of the HMA domain of Pikm are essential for the full binding to AVR-PikD (**Fig 8** and **9**).

**Fig 9.**
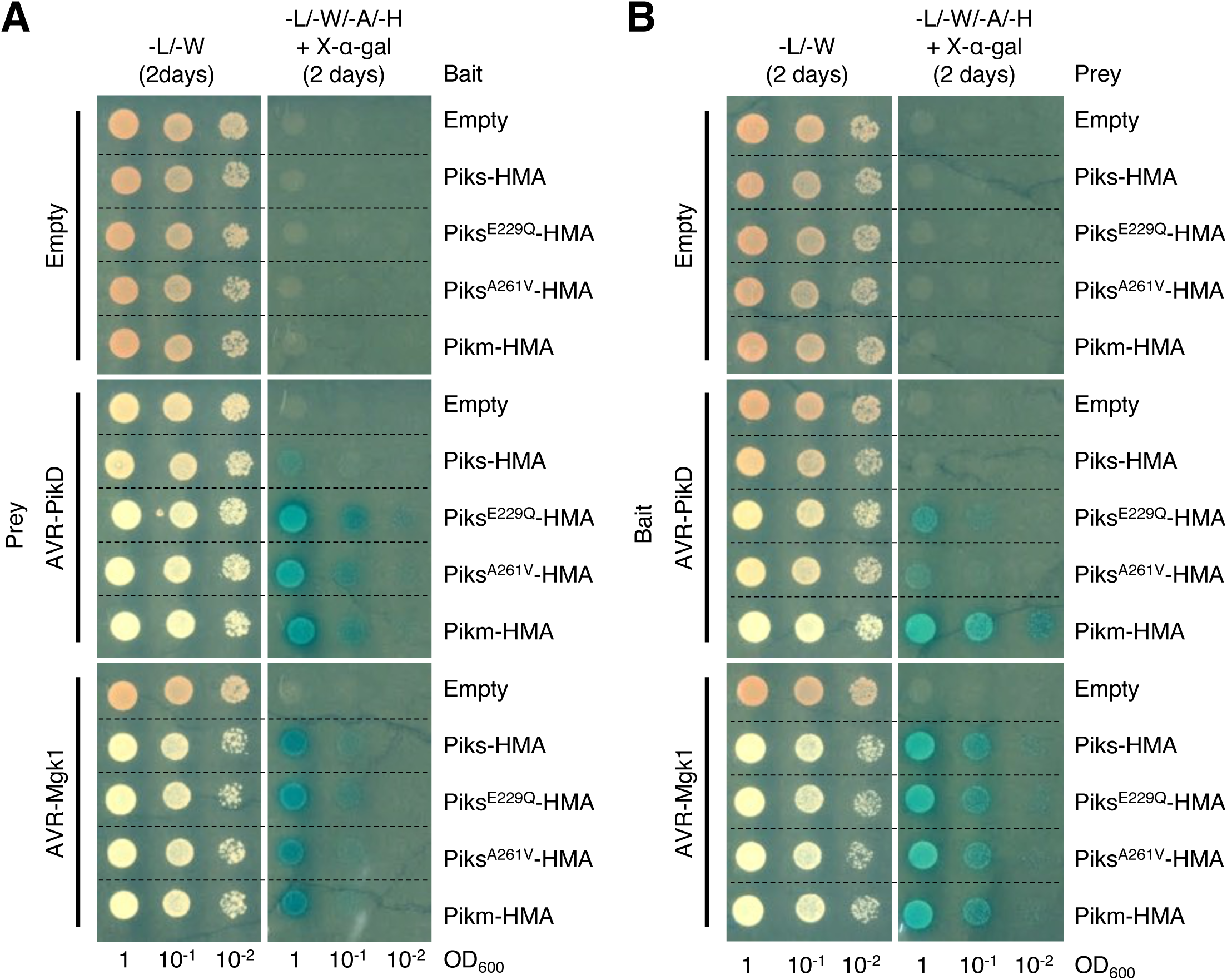
Yeast two-hybrid assay shows that the polymorphisms (E229Q and A261V) between Piks-HMA and Pikm-HMA quantitatively affect their binding to AVR-PikD. (A) HA-tagged AVRs as prey and Myc-tagged HMA domains as bait. (B) Myc-tagged AVRs as bait and HA-tagged HMA domains as prey. Empty vector was used as a negative control. Left side: basal medium lacking leucine (L) and tryptophan (W) for growth control. Right side: basal medium lacking leucine (L), tryptophan (W), adenine (A), and histidine (H) and containing X-α-gal for selection.

## Discussion

In this study, we revealed a gene-for-gene interaction between the well-studied rice *Pik* resistance gene and *M. oryzae* effector genes. We discovered that the *Pik* allele *Piks* encodes a protein that detects the *M. oryzae* effector AVR-Mgk1, a secreted protein that does not belong to the AVR-Pik effector family. Piks specifically detects and responds to AVR-Mgk1, but other Pik proteins detects AVR-Mgk1 and AVR-Pik, indicating a complex network of gene-for-gene interactions (**Fig 10**, **S7 Table**). The response of Pik-1 to AVR-Mgk1 was previously overlooked; this illustrates the challenge of unravelling complex gene-for-gene interactions using classical genetic approaches and highlights the dynamic nature of the co-evolution between an NLR integrated domain and multiple families of effector proteins. As illustrated in **Fig 10**, our understanding of the interactions between *M. oryzae AVR* effectors and rice disease resistance genes has gone beyond Flor’s single gene-for-gene model and involves network-type complexity at multiple levels [2–5].

**Fig 10.**
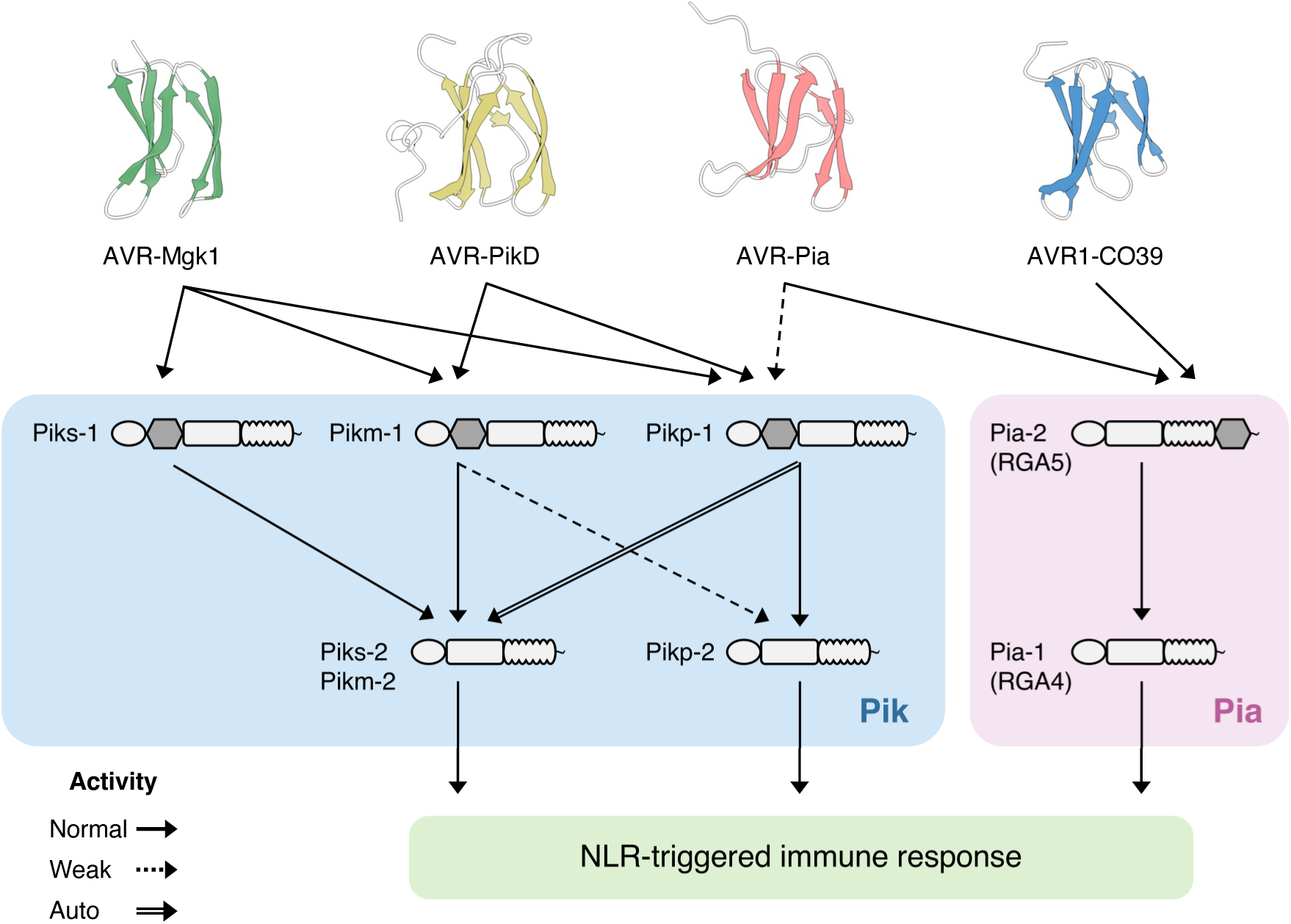
Beyond the gene-for-gene model: complex interactions between MAX effectors and rice NLR pairs. The NLR pairs Pik (Pik-1/Pik-2) and Pia (Pia-2/Pia-1, also known as RGA5/RGA4) have an integrated HMA domain (grey) in their sensor NLRs (Pik-1 and Pia-2). The Pia-2 HMA domain binds the sequence-unrelated MAX effectors AVR-Pia and AVR1-CO39 [69]. The Pikp-1 HMA domain weakly binds AVR-Pia, while that of Pikm-1 cannot [116]. The AVR-Mgk1 effector is detected by several Pik proteins, including Piks, which does not respond to AVR-PikD. Complex interactions also occur between sensor and helper NLRs forming homo- and hetero-complexes [98, 118]. An allelic mismatch of a receptor pair leads to autoimmunity (Pikp-1/Pikm-2) or reduced response (Pikm-1/Pikp-2) due to allelic specialization [119].The structures of AVR-Mgk1 predicted by AlphaFold2 [108], AVR-PikD (PDB ID: 6FU9 chain B) [71], AVR-Pia (PDB ID: 6Q76 chain B) [116], and AVR1-CO39 (PDB ID: 5ZNG chain C) [75] were visualised by ChimeraX [120].

### Why was the response of Pik-1 to AVR-Mgk1 previously overlooked?

Despite its recognition by multiple Pik proteins, AVR-Mgk1 had not been discovered by previous studies. This is mainly because *AVR-Mgk1* sequences are rare among the available *M. oryzae* genome sequences (**S6 Table**). In addition, the mini-chromosome encoding *AVR-Mgk1* appears to be absent from many isolates, and thus has no homologous chromosome sequence to recombine with. Our TH3o x O23 cross resulted in constantly similar significant *p*-values in the genetic association analysis (**Fig 4C**). The mini-chromosome is also affected by segregation distortion, resulting in a lower-than-expected frequency of *AVR-Mgk1* inheritance in the F_1_ progeny (**S5A Fig**). Lastly, the mini-chromosome of the O23 isolate carries two distinct *AVR* genes, *AVR-Mgk1* (two copies) and *AVR-PikD_O23*, which are both recognized by a single *Pik-1* resistance gene (**Fig 4C**, **S10 Fig**). *AVR-Mgk1* and *AVR-PikD_O23* mask each other’s activities and are tightly linked on the mini-chromosome, which is unfavourable for identification using classical genetic approaches.

Another challenge for identifying *AVR-Mgk1* was that the rice *Pish* locus, which confers resistance to O23 and TH3o, is also present in the rice cultivar Hitomebore (which contains *Piks*) (**Fig 2B** and **2C**). Thus, this network of gene-for-gene interactions was complicated by mutually masking *AVR* genes as well as by stacked and paired rice resistance genes. Disentangling the overlapping contributions of these resistance loci required rice RILs lacking the *Pish* locus (**Fig 2**). Therefore, unravelling complex networks of gene-for-gene interactions requires multiple-organism genetic approaches. This also demonstrates that to fully exploit genetic resistance, we need to go beyond the ‘blind’ approach of breeding and deploying *R* genes in agricultural crops without knowledge of the identity and population structure of the *AVR* genes encoding the effectors they are potentially sensing.

### The *AVR-Mgk1* genes are flanked by retrotransposon sequences

We observed deletion of *AVR-Mgk1* genes in one out of 144 sexual recombinants in just one generation. This event was mediated by the tandemly duplicated *Inago2* retrotransposon sequences that flank the *AVR-Mgk1* genes (**Fig 5A**). We hypothesize that this type of repeat sequence-mediated deletion of *AVR* genes might occur frequently in nature. The *M. oryzae* effector gene *AVR-Pita*, which occurs on the same mini-chromosome as *AVR-Mgk1* and *AVR-PikD_O23*, is also flanked by the solo long terminal repeats (solo-LTRs) of the retrotransposons *Inago1* and *Inago2* near the telomeric end of the chromosome [47] opposite of *AVR-Mgk1* and *AVR-PikD_O23* (**Fig 4C**). Chuma et al. proposed that the linkage of *AVR*-*Pita* to retrotransposons is associated with translocation between different *M. oryzae* isolates, and therefore, may facilitate horizontal gene transfer and recovery, particularly in asexual lineages [47]. This effector gene– retrotransposon linkage could enable persistence of the effector gene in the fungal population despite repeated deletions, and is a potential mechanism underpinning the two-speed genome concept [121–123]. In the case of *AVR-Mgk1*, *Inago2* and dense solo-LTRs located between the two *AVR-Mgk1* copies (**Fig 5A**) appear to contribute to the effector gene’s genetic instability and may explain its low frequency in *M. oryzae* populations.

### AVR-Mgk1 is predicted to adopt a MAX fold structure

Despite the primary sequence dissimilarity between AVR-Mgk1 and AVR-Pik, AlphaFold2 [108] predicted that AVR-Mgk1 adopts a MAX fold structure (**Fig 6C**) similar to AVR-Pik and several other *M. oryzae* effectors [27–30,35,124]. However, the region that includes the highly polymorphic residues of AVR-Pik effectors, which determine their binding to the HMA domain of Pik-1 and are modulated by arms race co-evolution [70,71,96], differs structurally in AVR-Mgk1 (**Fig 6C**–**E**). This suggests that the HMA domain may bind AVR-Mgk1 at different interacting residues (or a subset of different interacting residues) from AVR-Pik as demonstrated for other MAX effectors [72,74,75,116]. This is supported by the observation that the Piks polymorphisms, which alter binding to AVR-PikD, do not affect the interaction with AVR-Mgk1 (**Fig 8** and **9**). It is remarkable that *M. oryzae* effectors may have evolved to bind the HMA domain through multiple interfaces, which necessitates additional structural studies of effector–HMA complexes.

### Identification of AVR-Mgk1 highlights flexible and complex host–pathogen recognition by an integrated domain

The identification of AVR-Mgk1 expands our understanding of the interaction between the integrated HMA domains of rice NLR receptors and MAX effectors (**Fig 10**). Pik proteins Pikm, Pik*, and Pikp detect and bind AVR-Mgk1 and AVR-PikD via the Pik-1 integrated HMA domain (**S9** and **S12**–**16 Figs**). The recognition of multiple MAX effectors by an NLR receptor was reported in the rice NLR pair Pia [69]. The sensor NLR Pia-2 (RGA5) also contains the HMA domain, which binds the sequence-unrelated MAX effectors AVR-Pia and AVR1-CO39 [69,74,75]. The presence of the HMA domain in Pik proteins also enables Pikp to weakly respond to AVR-Pia, while this response is not observed with the combination of Pikm and AVR-Pia [116]. These reports indicate that an integrated domain can flexibly recognize multiple pathogen effectors. Our findings further extend the knowledge of HMA-mediated MAX effector recognition in that the recognition specificity of AVR-Mgk1 is different from that of previously identified MAX effectors, such as AVR-PikD, AVR-Pia, and AVR1-CO39 (**Fig 10**). The AVR-Mgk1- and AVR-PikD-interacting residues (or a subset of interacting residues) of the Pik HMA domain likely differ (**Fig 6**, **8**, and **9**). These different modes of interactions would enable an HMA domain to target multiple effectors, and therefore contribute to a broad recognition spectrum for pathogen effectors.

In the interactions between Pik proteins and AVR-Pik effectors, only a few polymorphisms dynamically change the recognition spectrum and determine the recognition specificity [70–73,96]. Here, we demonstrated that Piks binds and responds to AVR-Mgk1, but not to AVR-PikD (**Fig 7**). This unique recognition spectrum of Piks among other Pik family proteins (**Fig 10**) is caused by two amino acid changes (E229Q and A261V) relative to its quasi-identical protein Pikm (**Fig 8** and **9**). We could not unambiguously reconstruct the ancestral state and evolutionary trajectory of these two key polymorphisms because they are recurrently polymorphic among Pik-1 proteins. However, considering that these polymorphisms between Piks-1 and Pikm-1 have arisen among cultivated rice, Piks-1 may have lost the capacity to respond to AVR-PikD as a trade-off between Pik immunity and rice yield, as reported for another rice resistance gene, *Pigm* [125].

Collectively, our findings imply the potential of integrated HMA domains to flexibly recognize pathogen effectors. In parallel, arms race co-evolution with *M. oryzae* and agricultural selection generate HMA domain variants with different recognition specificities, which results in a network of tangled gene-for-gene interactions between integrated HMA domains and MAX effectors (**Fig 10**). HMA–effector interactions can be a model to understand the flexible and complex mechanisms of host–pathogen recognition established during their co-evolution.

## Materials and Methods

### *Magnaporthe oryzae* isolates O23 and TH3o and their genetic cross

The *Magnaporthe oryzae* isolates used in this study were imported to Japan with permission from the Ministry of Agriculture, Forest and Fishery (MAFF), Japan and are maintained at Iwate Biotechnology Research Center under the license numbers “TH3: MAFF directive 12 yokoshoku 1139” and “O23: MAFF directive 51 yokoshoku 2502”. Genetic crosses of the *M. oryzae* isolates TH3o (subculture of TH3) and O23 (O-23IN [PO12-7301-2]) [47] were performed as previously described [126]. Briefly, perithecia were formed at the intersection of mycelial colonies of TH3o and O23 on oatmeal agar medium (20 g oatmeal, 10 g agar, and 2.5 g sucrose in 500 ml water) in a Petri dish during 3–4 weeks of incubation at 22°C under continuous fluorescent illumination. Mature perithecia were crushed to release asci, which were transferred to a water agar medium (10 g of agar in 500 ml of water) with a pipette. Each ascus was separated with a fine sterilized glass needle under a micromanipulator. After 24 h incubation, germinated asci were transferred to potato dextrose agar (PDA) slants. After two weeks incubation, the resulting mycelial colonies were used for spore induction, and the spore solution was diluted and spread on PDA medium. After a 1-week incubation, a mycelial colony derived from a single spore was transferred and used as an F_1_ progeny of TH3o and O23. For long-term storage, the F_1_ progeny was grown on sterilized barley (*Hordeum vulgare*) seeds in vials at 25°C for one month and kept in a case with silica gel at 10°C.

### *M. oryzae* infection assays

For infection assay, rice plants one month after sowing were used. *M. oryzae* isolates TH3o, O23, and their F_1_ progeny were grown on oatmeal agar medium [40% oatmeal (w/v), 5% sucrose (w/v), and 20% agar (w/v)] for two weeks at 25°C. Then aerial mycelia were washed off by rubbing mycelial surfaces with plastic tube, and the colonies were incubated under black light (FL15BLB; Toshiba) for 3 to 5 days to induce conidiation. Resulting conidia were suspended in distilled water, and adjusted to the concentration of 5 × 10^5^ spores per ml. The conidial suspension was inoculated onto the press-injured sites on rice leaves. The inoculated plants were incubated under dark at 28°C for 20 h, and then transferred to a growth chamber at 28°C with a 16-h light/8-h dark photoperiod. Disease lesions were photographed 10–12 days after inoculation. The vertical lesion length was measured. The lesion sizes were plotted by the “barplot” and “swarmplot” functions in the “seaborn” python library v0.11.2. The confidence interval was calculated by the default parameters of the “barplot” function. Two-sided Welch’s t-test was conducted by the “ttest_ind” function in the “SciPy” python library v1.7.2 with the option “equal_var: False, alternative: two-sided”.

### Sequencing of rice cultivars Hitomebore and Moukoto and RILs derived from their cross

We re-sequenced rice (*Oryza sativa*) lines Hitomebore and Moukoto and 249 RILs from their cross. First, genomic DNA was extracted from fresh leaves using Agencourt Chloropure Kit (Beckman Coulter, Inc, CA, USA). Then, DNA was quantified using Invitrogen Quant-iT PicoGreen dsDNA Assay Kits (Thermo Fisher Scientific, MA, USA). For Hitomebore and Moukoto, library construction was performed using TruSeq DNA PCR-Free Library Prep Kit (Illumina, CA, USA). These two libraries were sequenced using the Illumina NextSeq, HiSeq, and MiSeq platforms (Illumina, CA, USA) for 75-bp, 150-bp, and 250/300-bp paired-end reads, respectively (**S1 Table**). For the 249 RILs, library construction was performed using house-made sequencing adapters and indices. These libraries were sequenced using the Illumina NextSeq platform for 75 bp paired-end reads (**S1 Table**). First, we removed adapter sequences using FaQCs v2.08 [127]. Then, we used PRINSEQ lite v0.20.4 [128] to remove low-quality bases with the option “-trim_left 5-trim_qual_right 20 -min_qual_mean 20 -min_len 50.” In addition, 300-bp reads were trimmed to 200 bp by adding an option “-trim_to_len 200.”

### SNP calling for the rice RIL population

The quality-trimmed short reads of the two parents and 249 RILs were aligned to the reference genome of Os-Nipponbare-Reference-IRGSP-1.0 [129] using bwa mem command in BWA v0.7.17 [130] with default parameters. Using SAMtools v1.10 [131], duplicated reads were marked, and the alignments were sorted in positional order. These BAM files were subjected to variant calling. First, we performed valiant calling for the parent cultivars Hitomebore and Moukoto according to the “GATK Best Practices for Germline short variant discovery” [132] (https://gatk.broadinstitute.org/), which contains a BQSR step, two variant calling steps with HaplotypeCaller in GVCF mode and GenotypeGVCFs commands, and a filter valiant step with VariantFiltration command with the option “QD < 2.0 || FS > 60.0 || MQ < 40.0 || MQRankSum < -12.5 || ReadPosRankSum < -8.0 || SOR > 4.0.” In the resulting VCF file, we only retained biallelic SNPs where: 1) both parental cultivars had homozygous alleles, 2) the genotypes were different between Hitomebore and Moukoto, and 3) both parental cultivars had a depth (DP) of eight or higher. As a result, 156,503 SNP markers were extracted, and the position of these SNPs was converted to a bed file (position.bed) using the BCFtools query command. For SNP genotyping of the 249 RILs, the VCF file was generated as follows: 1) BCFtools v1.10.2 [133] mpileup command with the option “-t DP,AD,SP -A -B -Q 18 -C 50 -uv -l position.bed”; 2) BCFtools call command with the option “-P 0 -A -c -f GQ”; 3) BCFtools filter command with the option “-v snps -i ’INFO/MQ>=0 & INFO/MQ0F<=1 & AVG(GQ)>=0”; 4) BCFtools norm command with the option “-m+both. ” Finally, we imputed the variants based on Hitomebore and Moukoto genotypes using LB-impute [134].

### *De novo* assembly of the Hitomebore genome

To reconstruct the *Pish* and *Pik* regions in Hitomebore, we performed a *de novo* assembly using Nanopore long reads and Illumina short reads. To extract high-molecular-weight DNA from leaf tissue for nanopore sequencing, we used the NucleoBond high-molecular-weight DNA kit (MACHEREY-NAGEL, Germany). After DNA extraction, low-molecular-weight DNA was eliminated using the Short Read Eliminator Kit XL (Circulomics, Inc, MD, USA). Then, following the manufacturer’s instructions, sequencing was performed using Nanopore PromethION (Oxford Nanopore Technologies [ONT], UK). First, base-calling of the Nanopore long reads was performed for FAST5 files using Guppy 3.4.5 (ONT, UK), converted to FASTQ format (**S1 Table**). The lambda phage genome was removed from the generated raw reads with NanoLyse v1.1.0 [135]. We then trimmed the first 50 bp of each read and filtered out reads with an average read quality score of less than seven and reads shorter than 3,000 bases with NanoFilt v2.7.1 [135]. Next, the Nanopore long reads were assembled using NECAT v0.0.1 [136] setting the genome size to 380 Mbp. To further improve the accuracy of assembly, Racon v1.4.20 [137] was used twice for error correction, and Medaka v1.4.1 (https://github.com/nanoporetech/medaka) was subsequently used to correct mis-assembly. Following this, two rounds of consensus correction were performed using bwa-mem v0.7.17 [130] and HyPo v1.0.3 [138] with Illumina short reads. We subsequently removed haplotigs using purge-haplotigs v1.1.1 [139], resulting in a 374.8 Mbp *de novo* assembly comprising 77 contigs. This assembly was further scaffolded with RagTag v1.1.0 [140], with some manual corrections, using the Os-Nipponbare-Reference-IRGSP-1.0 as a reference genome. The resulting Hitomebore genome sequence was deposited on Zenodo (https://doi.org/10.5281/zenodo.7317319).

### RNAi-mediated knockdown of *Piks-1* and *Piks-2* in rice

To prepare *Piks-1* and *Piks-2* knockdown vectors, the cDNA fragments Piks-1A (nt 618–1011) and Piks-1B (nt 1132–1651) for *Piks-1*, and Piks-2A (nt 121–524) and Piks-2B (nt 2317–2726) for *Piks-2* were amplified using primer sets (KF852f/KF853r, KF854f/KF855r, KF848f/KF849r, and KF801f/KF802r, respectively, **S8 Table**). The resulting PCR products were cloned into the Gateway vector pENTR-D-TOPO (Invitrogen, Carlsbad, CA, USA), and transferred into the pANDA vector [141] using LR clonase (Invitrogen), resulting in pANDA-Piks-1A, pANDA-Piks-1B, pANDA-Piks-2A, and pANDA-Piks-2B. Plasmids were transformed into *Agrobacterium tumefaciens* (EHA105) and used for stable transformation of rice RIL #58 (*Piks* +) by *Agrobacterium-*mediated transformation. Transformation and regeneration of rice plants were performed according to Hiei et al. [142].

To determine *Piks-1* and *Piks-2* expression in the transgenic lines, reverse transcription quantitative PCR (RT-qPCR) was performed. Total RNA was isolated from transformant leaves using the Qiagen RNeasy plant mini kit (Qiagen, Venlo, the Netherlands). cDNA was synthesized with the ReverTra Ace kit (TOYOBO, http://www.toyobo.co.jp) and used as a template for quantitative PCR (qPCR) using primer sets (YS29f/YS30r for *Piks-1*, YS35f/YS36r for *Piks-2*, Actin-RTf/Actin-RTr for rice *Actin*, **S8 Table**). qPCR was performed using the Luna Universal qPCR Master Mix (New England Biolabs Japan, Tokyo, Japan) on a QuantStudio 3 Real-Time PCR System (Thermo Fisher Scientific, MA, USA). The relative expression levels of *Piks*-1 and *Piks*-2 were calculated via normalization with rice *Actin*. The relative expression levels were plotted by the “barplot” and “swarmplot” functions in the “seaborn” python library v0.11.2. The confidence interval was calculated by the default parameters of the “barplot” function. A two-sided Welch’s t-test was conducted by the “ttest_ind” function in the “SciPy” python library v1.7.2 with the option “equal_var: False, alternative: two-sided”.

### Phylogenetic analysis of Pik alleles

The sequences of *Pik-1* (*Pikh-1* [AET36549.1], *Pikp-1* [ADV58352.1], *Pik*-1* [ADZ48537.1], *Pikm-1* [AB462324.1], and *Piks-1* [AET36547.1]) and *Pik-2* (*Pikh-2* [AET36550.1], *Pikp-2* [ADV58351.1], *Pik*-2* [ADZ48538.1], *Pikm-2* [AB462325.1], and *Piks-2* [AET36548.1]) were aligned using MAFFT v7.490 [143] with the option “--globalpair --maxiterate 1000”. The phylogenetic trees of *Piks-1* and *Piks-2* were separately drawn based on nucleotide sequences with IQ-TREE v2.0.3 [144] using 1,000 ultrafast bootstrap replicates [145]. The models for reconstructing trees were automatically selected by ModelFinder [146] in IQ-TREE. ModelFinder selected “HKY+F” for *Pik-1* and “F81+F” for *Pik-2* as the best-fit models according to the Bayesian information criterion (BIC). Finally, the midpoint rooted trees were drawn with FigTree v1.4.4 (http://tree.bio.ed.ac.uk/software/figtree/).

### Sequencing of *M. oryzae* isolates O23 and TH3o and their F_1_ progeny

For long-read sequencing, O23 and d44a genomic DNA was extracted from liquid-cultured aerial hyphae using the NucleoBond high-molecular-weight DNA kit (MACHEREY-NAGEL, Germany). The genomic DNA was processed through the short-read eliminator kit XL (Circulomics). The filtered genomic DNA (2 µg) was used to construct a library for Nanopore sequencing using the ligation sequencing kit SQK-LSK109 (ONT, UK). Sequencing was performed using the MinION system with a FLO-MIN106D (R9.4) flow cell (ONT, UK).

TH3o genomic DNA was extracted using the cetyl trimethyl ammonium bromide (CTAB) method. The extracted DNA was purified using Genomic-tip (Qiagen, Germany) according to the manufacturer’s protocol. Sequencing was performed by Macrogen, Inc., Seoul, Korea, using the PacBio RS II sequencer (Pacific Biosciences of California, Inc., Menlo Park, CA, USA).

For short-read sequencing of O23, TH3o, and their F_1_ progeny, genomic DNA was extracted from aerial hyphae using the NucleoSpin Plant II Kit (Macherey Nagel). Libraries for paired-end short reads were constructed using an Illumina TruSeq DNA LT Sample Prep Kit (Illumina, CA, USA). The paired-end library was sequenced by the Illumina NextSeq platform (Illumina, CA, USA). We also sequenced O23 genomic DNA using the MiSeq platform to polish the *de novo* O23 assembly.

The adapters of short-reads were trimmed by FaQCs v2.08 [127]. In this step, we also filtered the reads and discarded reads shorter than 50 bases and those with an average read quality below 20.

### *De novo* assembly of O23, TH3o, and d44a genomes

First, base-calling of the Nanopore long reads was performed for FAST5 files of O23 and d44a with Guppy 3.4.4 (ONT, UK). The lambda phage genome was removed from the generated raw reads with NanoLyse v1.1.0 [135]. We then trimmed the first 50 bp of each read and filtered out reads with an average read quality score of less than 7 and reads shorter than 3,000 bases with NanoFilt v2.7.1 [135]. The quality-trimmed Nanopore long reads of O23 and d44a were assembled with NECAT v0.0.1 [136] setting the genome size to 42 Mbp. The assembled contigs were then polished with medaka v0.12.1 (https://github.com/nanoporetech/medaka) and with Hypo v1.0.3 [138]. In Hypo, we used MiSeq and NextSeq short-reads for O23 and d44a, respectively, in addition to quality-trimmed Nanopore long reads.

For the *de novo* assembly of TH3o, we trimmed the first 50 bp of each read and filtered out reads with an average read quality score of less than 7 and reads shorter than 2,000 bases with NanoFilt v2.7.1 [135]. The quality-trimmed PacBio long reads of TH3o were assembled with MECAT v2 [147] setting the genome size to 42 Mbp. The assembled contigs were polished with Hypo v1.0.3 [138] using NextSeq short-reads and PacBio long reads of TH3o.

To evaluate the completeness of the gene set in the assembled contigs, we applied BUSCO analysis v3.1.0 [99]. For BUSCO analysis, we set “genome” as the assessment mode, and *Magnaporthe grisea* was used as the species in AUGUSTUS [148]. *Sordariomyceta* odb9 was used as the dataset.

The genome sequences of the *M. oryzae* isolates 70-15 (MG8 genome assembly in https://fungi.ensembl.org/Magnaporthe_oryzae/Info/Index) [100], O23, TH3o, and d44a were compared by dot plot analysis of D-GENIES [149]. The chromosome sequences of O23 and d44a were numbered and ordered based on those of 70-15.

### Variant calling for the *M. oryzae* F_1_ progeny derived from a cross between O23 and TH3o

Quality-trimmed short-reads were aligned to the O23 reference genome using the bwa mem command in BWA v0.7.17 with default parameters [130]. Using SAMtools v1.10 [131], duplicated reads were marked and the alignments were sorted to positional order. Only properly paired and uniquely mapped reads were retained using SAMtools [131]. For SNP markers on core chromosomes (chromosomes 1–7), the VCF file was generated as follows: 1) BCFtools v1.10.2 [133] mpileup command with the option “-a AD,ADF,ADR -B -q 40 -Q 18 -C 50”; 2) BCFtools call command with the option “-vm -f GQ,GP --ploidy 1”; 3) BCFtools filter command with the option “-i “INFO/MQ>=40”. In the VCF file, biallelic SNPs were retained only where: 1) O23 had the same genotype as the O23 reference genome, 2) both parental isolates, O23 and TH3o, had a depth (DP) of four or higher, 3) the average genotype quality (GQ) across all the samples was 100 or higher, 4) the number of missing genotypes among the 144 F_1_ progeny was less than 15, and 5) the allele frequency was between 0.05 and 0.95. As a result, 7,867 SNP markers were extracted from the core chromosomes. For presence/absence markers on the remaining contigs, we selected candidate presence/absence regions on the parental genomes, O23 and TH3o. First, the BCFtools mpileup command was used only for the BAM files of O23 and TH3o with the option “-a DP -B -q 40 -Q 18 -C 50”. Second, BCFtools view command was used with the option “-g miss -V indels” to extract the positions where either O23 or TH3o was missing. Third, only the positions where O23 had a depth of eight or higher and TH3o had a depth of zero were retained. These positions were concatenated using the bedtools v2.29.2 [150] merge command with the option “-d 10”. Only candidate regions larger than or equal to 50 bp were retained. Using the SAMtools bedcov command with the option “-Q 0”, the number of alignments of each F_1_ progeny on these candidate regions was counted. If an F_1_ progeny had at least one alignment on a candidate region, the F_1_ progeny was considered to have a presence-type marker for that region. On the other hand, if an F_1_ progeny had no alignment on a candidate region, the F_1_ progeny was considered to have an absence-type maker for that region. Finally, only the presence/absence markers that 1) had an average depth of four or higher for O23 regions, and one or less for TH3o regions, and 2) had an allele frequency between 0.05 and 0.95 were retained. As a result, 265 presence/absence markers were extracted for the remaining contigs.

### Annotation of the O23 reference genome

The segregation distortion of each marker was tested by a two-sided binomial test (*p* = 0.5). O23-specific regions were annotated by aligning TH3o contigs to the O23 reference genome with Minimap2 [151] using the option “-x asm5”. Transposable elements were annotated by EDTA v1.9.0 [152] with the option “--anno 1 --species others --step all”. Coding sequences of the genome assembly version MG8 of the *M. oryzae* isolate 70-15 [100] and the library of transposable elements curated in Chuma et al. [47] were also provided as input to EDTA. We only retained the annotations from the provided transposable elements. LTRs of retrotransposons were also annotated by EDTA, independently. The genes on the O23 reference genome were annotated by aligning the coding sequences of the genome assembly version MG8 of 70-15 using Spaln2 v2.3.3 [153]. The sequence similarity of the mini-chromosome sequence O23_contig_1 was analysed against the O23 core chromosomes using Minimap2 [151] with the option “-x asm5”. We filtered out the alignments shorter than 1 kbp or with a mapping quality less than 40. Finally, these sequence similarities were plotted by Circos v0.69.8 (http://circos.ca/) including other genomic features. For gene density, the overlapped gene annotations were regarded as a single gene annotation. The plotted figure does not include contigs smaller than O23_contig_1.

### Association analysis between genetic markers and phenotype

The association between the genetic markers and the phenotype was evaluated using the R package rrBLUP [154]. To correct the threshold of *p*-values for multiple testing, false discovery rate was used for the rice RILs and *M. oryzae* F_1_ progeny. For false discovery rate, the “multipletests” function in the “statsmodels” python library was used with the option “method: fdr_bh, alpha: 0.05”.

### RNA-seq to identify *AVR-Mgk1*

Total RNA of TH3o and O23 was extracted at different stages (24 and 48 h) of barley infection using the SV Total RNA Isolation System (Promega, WI, USA). One microgram of total RNA was used to prepare each sequencing library with the NEBNext Ultra II Directional RNA library prep kit (New England Biolabs Japan, Tokyo, Japan) following the manufacturer’s protocol. The library was sequenced by paired-end mode using the Illumina Hiseq X platform (Illumina, CA, USA).

For quality control, the reads were filtered and reads shorter than 50 bases and those with an average read quality below 20 and trimmed poly(A) sequences were discarded with FaQCs v2.08 [127]. The quality-trimmed reads were aligned to the O23 reference genome with HISAT2 v2.1 [155] with the options “--no-mixed --no-discordant --dta”. BAM files were sorted and indexed with SAMtools v1.10 [131], and transcript alignments were assembled with StringTie v2.0 [156] separately for each BAM file.

### Transformation of *M. oryzae* isolate Sasa2 with *AVR-Mgk1* and *AVR-PikD_O23*

To construct the *pCB1531-pex22p-AVR-Mgk1* expression vector, *AVR-Mgk1* was amplified by PCR using primer sets XbaI_O23_48h.1149.1-F and BamHI_O23_48h.1149.1-R (**S8 Table**) from cDNA of *M. oryzae* O23-infected barley leaf material. The PCR product was digested with *Xba*I and *Bam*HI and ligated into the *pCB1531-pex22p-EGFP* vector [18] using the *Xba*I and *Bam*HI sites to be exchanged with EGFP tag. To construct the *pCB1531-pex22p-AVR-PikD’*(*AVR-Pik-D_O23*) expression vector, a 0.3-kb fragment containing *AVR-PikD’* (*AVR-Pik-D_O23*) was amplified by PCR using the primers Xba1_kozak_pex31_U1 [18] and KF792r (**S8 Table**) from *M. oryzae* O23 genomic DNA. The PCR product and *pCB1531-pex22p-EGFP* expression vector were digested with *Xba*I and *Eco*RI to ligate *AVR-PikD_O23* into the position of the EGFP tag, generating *pCB1531-pex22p-AVR-PikD’*(*AVR-Pik-D_O23*). The resulting vectors were used to transform *M. oryzae* Sasa2 following a previously described method [157].

To confirm *AVR-Mgk1* expression in infected rice leaves, Sasa2 transformants were punch inoculated on rice cultivar Moukoto. We reverse transcribed cDNA from RNA extracted from the infected rice leaves and amplified *AVR-Mgk1* via PCR using primer sets listed in **S8 Table**. Rice and *M. oryzae Actin* were used as controls.

### Protein sequence alignment between AVR-Mgk1 and AVR-PikD

NCBI BLAST (https://blast.ncbi.nlm.nih.gov/Blast.cgi) was used to align the AVR-Mgk1 and AVR-PikD protein sequences using the Needleman-Wunsch algorithm [158] for pairwise global alignment using default parameters.

### Clustering of putative *M. oryzae* AVR protein sequences using TRIBE-MCL

A dataset of the putative *M. oryzae* effector proteins [32] amended with AVR-Mgk1 was clustered by TRIBE-MCL [107] using “1e-10” for an E-value cut-off of BLASTP [159] and “1.4” for the inflation parameter “-I” in mcl. The other parameters were default. The sequence set used in this analysis was deposited in Zenodo (https://doi.org/10.5281/zenodo.7317319).

### AVR-Mgk1 structure prediction

The AVR-Mgk1 structure was predicted using AlphaFold2 [108]. The signal peptide sequence in AVR-Mgk1 was predicted by SignalP v6.0 (https://services.healthtech.dtu.dk/service.php?SignalP) [106]. The amino acid sequence without the signal peptide (Arg25-Trp85) was used as an input for AlphaFold2 [108], available on the Colab notebook. The best model generated by AlphaFold2 was visualised by ChimeraX v1.2.5 [120] together with the protein structures of AVR-PikD (PDB ID: 6FU9 chain B) [71], AVR-Pia (PDB ID: 6Q76 chain B) [116], and AVR1-CO39 (PDB ID: 5ZNG chain C) [75]. The protein structures of AVR-Mgk1 and AVR-PikD were aligned by structure-based alignment using TM-align (https://zhanggroup.org/TM-align) [109]. The AVR-Mgk1 structure predicted by AlphaFold2 is deposited on Zenodo (https://doi.org/10.5281/zenodo.7317319).

### BLAST search of AVR-Mgk1 to the NCBI database

To find sequences related to AVR-Mgk1, BLASTN and BLASTP searches were run against the non-redundant NCBI database. A BLASTN search was also run against the whole-genome shotgun contigs of *Magnaporthe* (taxid: 148303). For all analyses, default parameters were used.

### Assays for protein-protein interactions

For the yeast two-hybrid assay, In-Fusion HD Cloning Kit (Takara Bio USA) was used to insert the *AVR-Mgk1* fragment (Arg25-Trp85) into pGADT7 (prey) and pGBKT7 (bait). DNA sequences of the fragments of AVR-PikD (Glu22-Phe113) and the Pik HMA domains (Piks-HMA [Gly186-Asp264], Pikp-HMA [Gly186-Asp263], Pik*-HMA [Gly186-Asp264], Pikm-HMA [Gly186-Asp264], Piks^E229Q^-HMA [Gly186-

Asp264], and Piks^A261V^-HMA [Gly186-Asp264], defined in De la Concepcion et al. [71]) were ligated into pGADT7 and pGBKT7 as described previously [70]. The primer sets used for PCR amplification of the fragments are listed in **S8 Table**. Yeast two-hybrid assays were performed as described previously [70] using a basal medium lacking leucine (L), tryptophan (W), adenine (A), and histidine (H) and containing 5-Bromo-4-Chloro-3-Indolyl α-D-galactopyranoside (X-α-gal) (Clontech) to detect interactions. The basal medium also contained 10 mM 3-amino-1,2,4-triazole (3AT) (Sigma) for selection, except for **Fig 9**.

Co-IP experiments of transiently expressed proteins in *Nicotiana benthamiana* were performed as described previously [70]. The protein regions used in the co-IP experiment were the same as those used in the yeast two-hybrid assay. We used N-terminally tagged FLAG:AVR and HA:HMA. The lysates of AVRs and HMA domains were diluted to compare the results at the same concentration and mixed (1:4, 1:2, or 1:1 ratio) *in vitro* to assemble the protein complex. For co-IP of HA-tagged proteins, Anti-HA affinity gel (Sigma) was used, and proteins were eluted by using 0.25 mg/ml HA peptide (Roche). HA- and FLAG-tagged proteins were immunologically detected using HRP-conjugated anti-HA 3F10 (Roche) and anti-FLAG M2 (Sigma), respectively. The primer sets used in this experiment are listed in **S8 Table**.

### Hypersensitive response cell death assay in *N. benthamiana*

Transient gene expression in *N. benthamiana* was performed by agroinfiltration according to methods described by van der Hoorn et al. [160]. Briefly, *A. tumefaciens* strain GV3101 pMP90 carrying binary vectors was inoculated from glycerol stock in liquid LB supplemented with 30 µg/ml rifampicin, 20 µg/ml gentamycin, and 50 µg/ml kanamycin and grown overnight at 28°C with shaking until saturation. Cells were harvested by centrifugation at 2000 × g at room temperature for 5 min. Cells were resuspended in infiltration buffer (10 mM MgCl_2_, 10 mM MES-KOH pH 5.6, 200 µM acetosyringone) and diluted to the appropriate OD_600_ (**S9 Table** and also see [82, 161]) in the stated combinations and left to incubate in the dark for 2 hours at room temperature prior to infiltration into 5-week-old *N. benthamiana* leaves. Hypersensitive cell death phenotypes were scored from 0 to 6 according to the scale in Maqbool et al. [29].

## Data Availability Statement

All the sequence data used in this study was deposited at European Nucleotide Archive (ENA, https://www.ebi.ac.uk/ena/browser/home) and the DNA Data Bank of Japan (DDBJ, https://www.ddbj.nig.ac.jp/index-e.html) with the study accessions PRJEB53625 and PRJDB13864,

## Acknowledgements

We thank Hiroe Utsushi, Akiko Hirabuchi, Yukie Hiraka, and Mari Iwai at Iwate Biotechnology Research Center, Japan, for technical support, and Phil Robinson at The Sainsbury Laboratory, UK for photography. We also thank Jeff Ellis for his comments on the preprint.

---

**S1 Fig.**
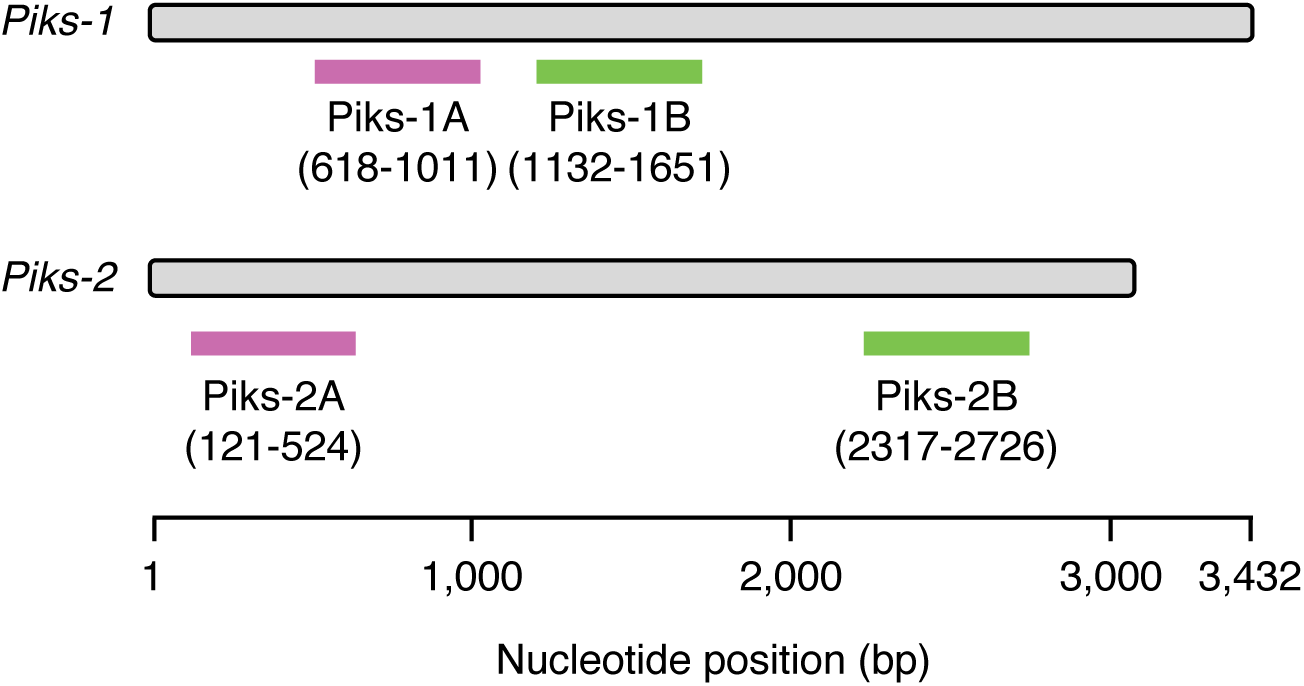
Schematic representations of the RNAi-mediated *Pik-1* and *Pik-2* knockdown experiment. For each *Pik* gene, we prepared two independent RNAi constructs targeting different regions on the gene (Piks-1A and Piks-1B for *Piks-1*, and Piks-2A and Piks-2B for *Piks-2*).

**S2 Fig.**
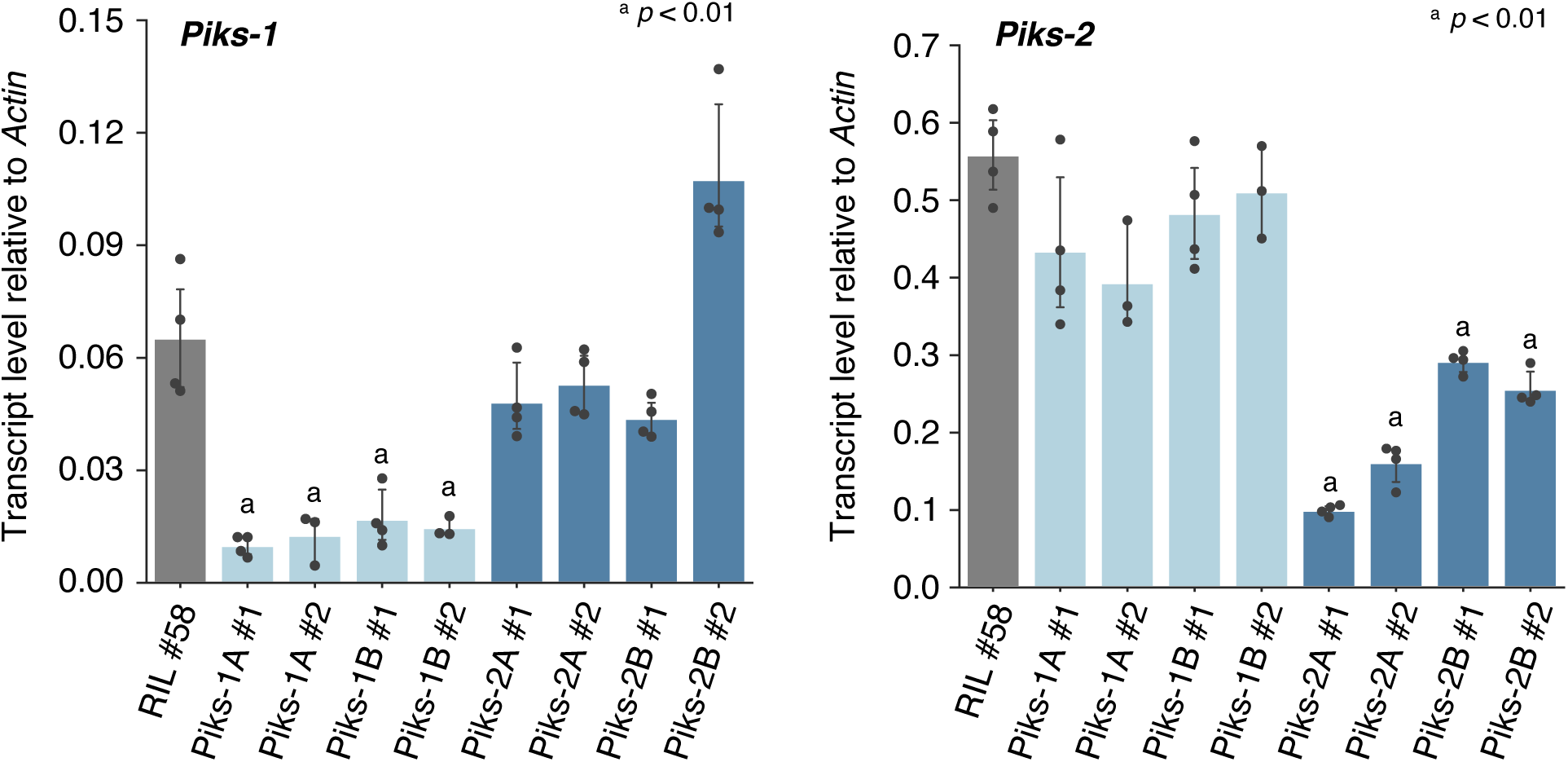
*Piks-1* and *Piks-2* expression in RNAi-mediated knockdown lines. We analyzed *Piks-1* and *Piks-2* expression in RNAi-mediated knockdown lines using RT-qPCR. RIL #58 (*Pish* -, *Piks* +) was used as the genetic background for the mutant lines. Rice *Actin* was used for normalization. ^a^ indicates statistically significant differences compared to RIL #58 (*p* < 0.01, two-sided Welch’s t-test). The data underlying this figure can be found in **S1 Data**.

**S3 Fig.**
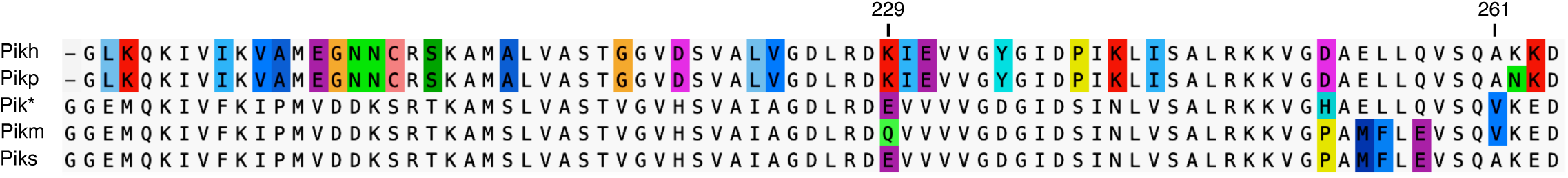
Amino acid sequence alignment of the HMA domains of Pik proteins. The sequences are visualized by the software AliView [162].

**S4 Fig.**
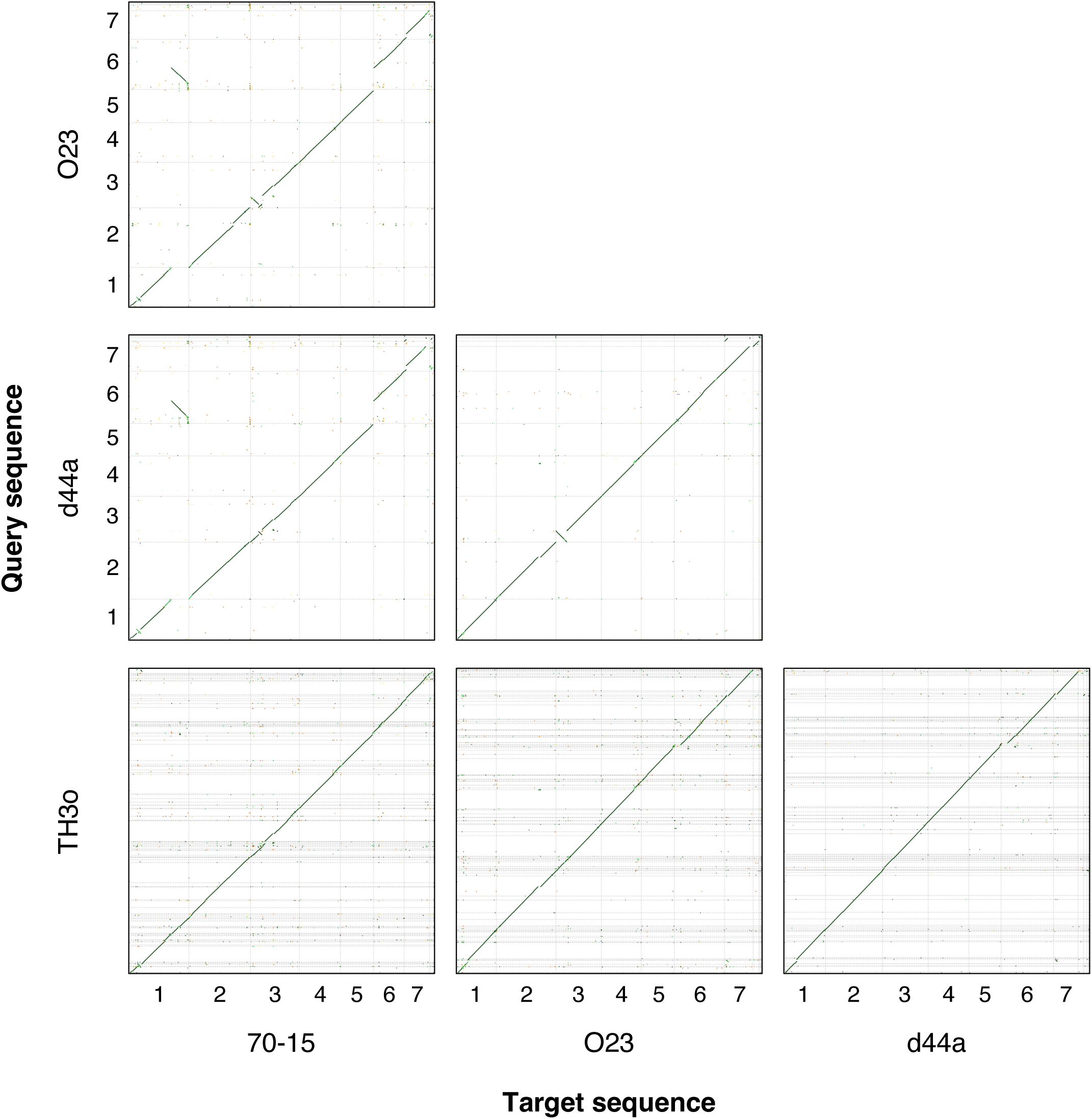
Pairwise dot plot analyses among the *de novo-*assembled genome sequences of *M. oryzae* isolates 70-15, O23, TH3o, and d44a. We compared the *de novo*-assembled genome sequences of O23, TH3o, and d44a with the previously assembled reference genome (MG8 genome assembly) of the isolate 70-15 [100], using D-GENIES [149]. The chromosome sequences of O23 and d44a are numbered and ordered based on those of 70-15. The data underlying this figure can be found in **S1 Data**.

**S5 Fig.**
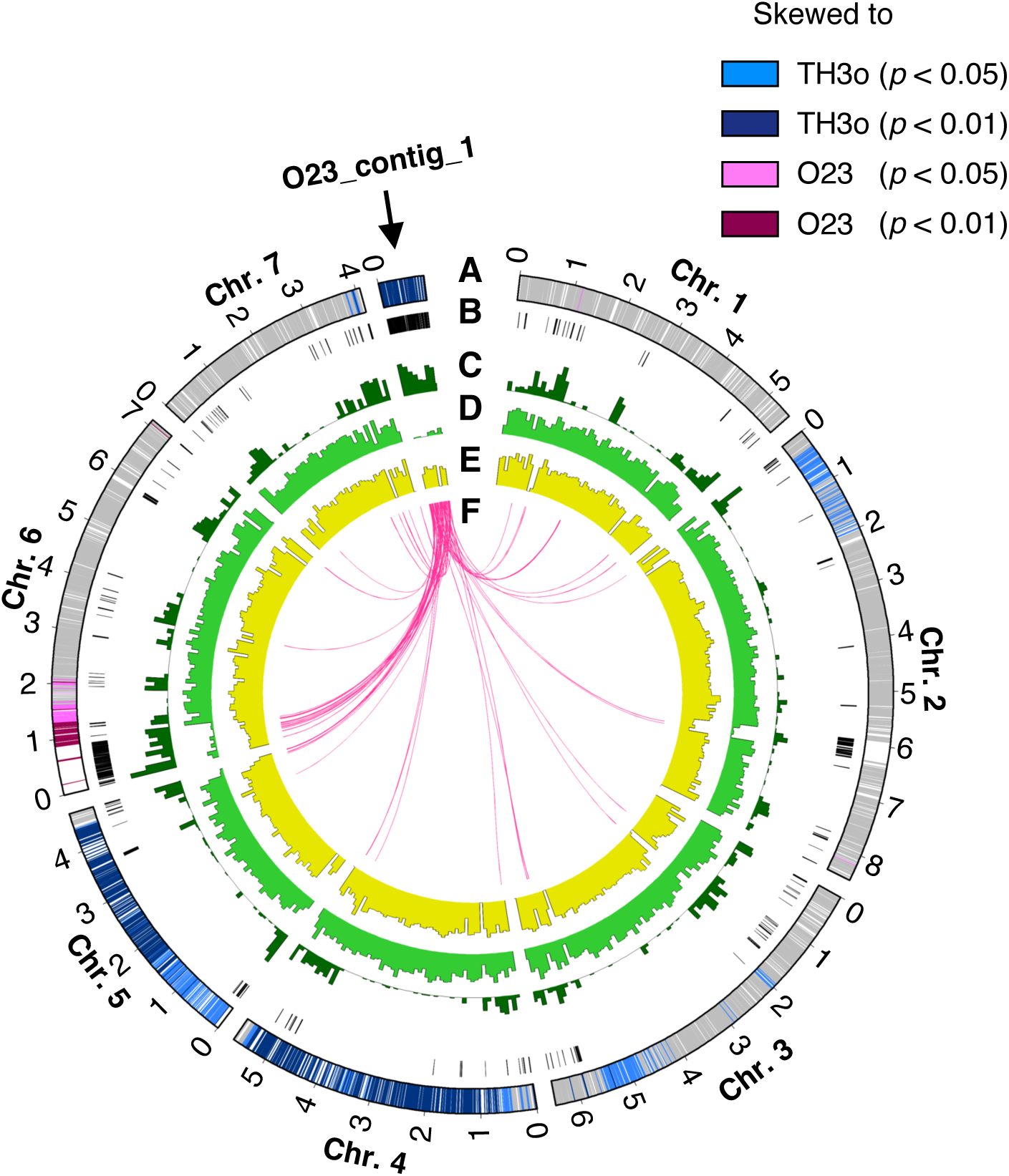
Genomic features of a *M. oryzae* mini-chromosome O23_contig_1. (A) Segregation distortion in TH3o x O23 F_1_ progeny. Grey markers are statistically neutral (*p* ≥ 0.05). (B) O23-specific regions (black) where TH3o contigs could not be aligned. (C) Density of transposable elements. (D) Gene density. (E) GC contents (0.45–0.55). (F) Sequence similarity between the O23 mini-chromosome and core chromosomes (chromosomes 1–7). The data underlying this figure can be found in **S1 Data**.

**S6 Fig.**
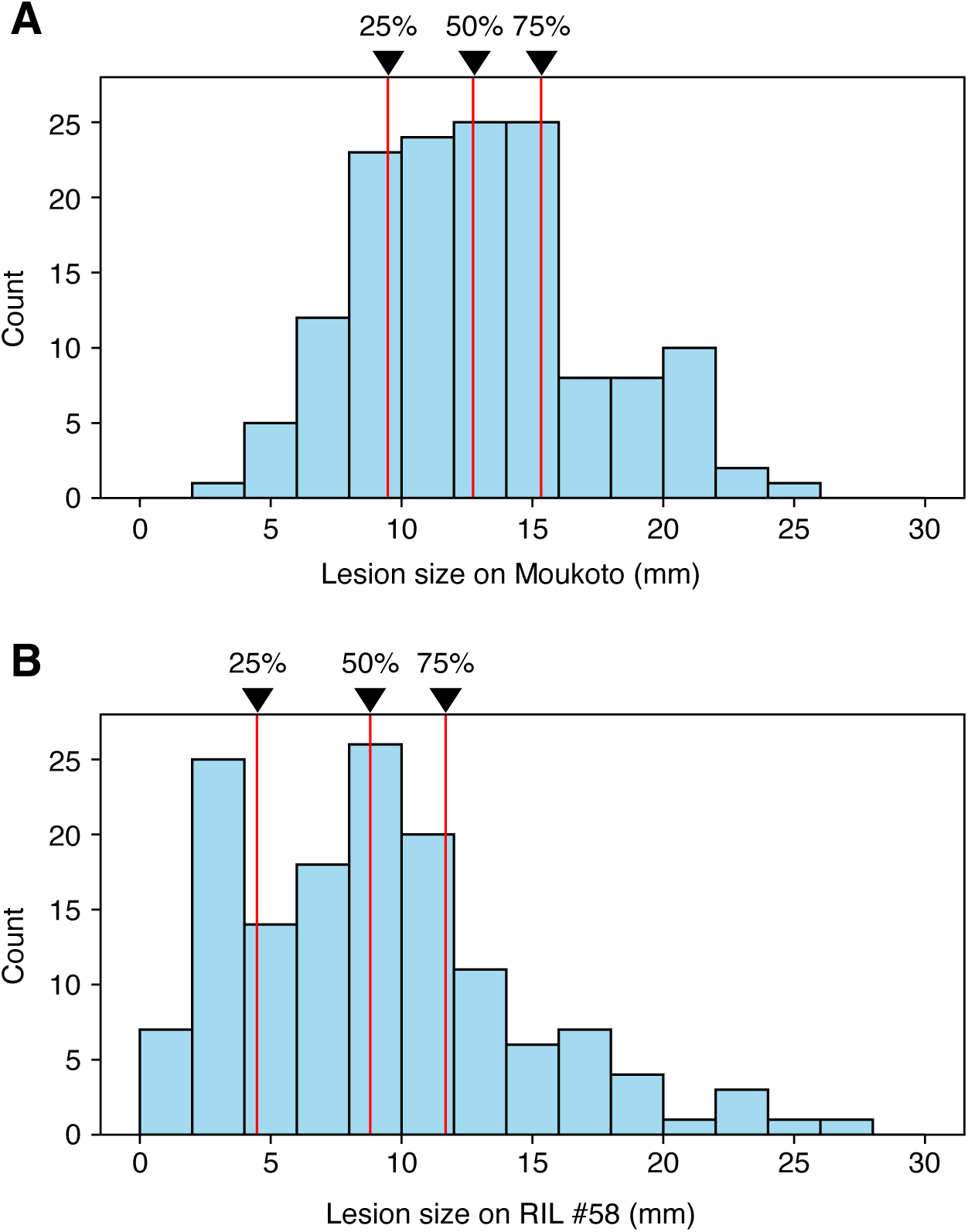
Histogram of the lesion size of F_1_ progeny in the rice inoculation assays to Moukoto and RIL #58. (A) Lesion size (mm) of F_1_ progeny on Moukoto. (B) Lesion size (mm) of F_1_ progeny on RIL #58. The percentiles (25%, 50%, and 75%) are indicated by the red lines. The data underlying this figure can be found in **S1 Data**.

**S7 Fig.**
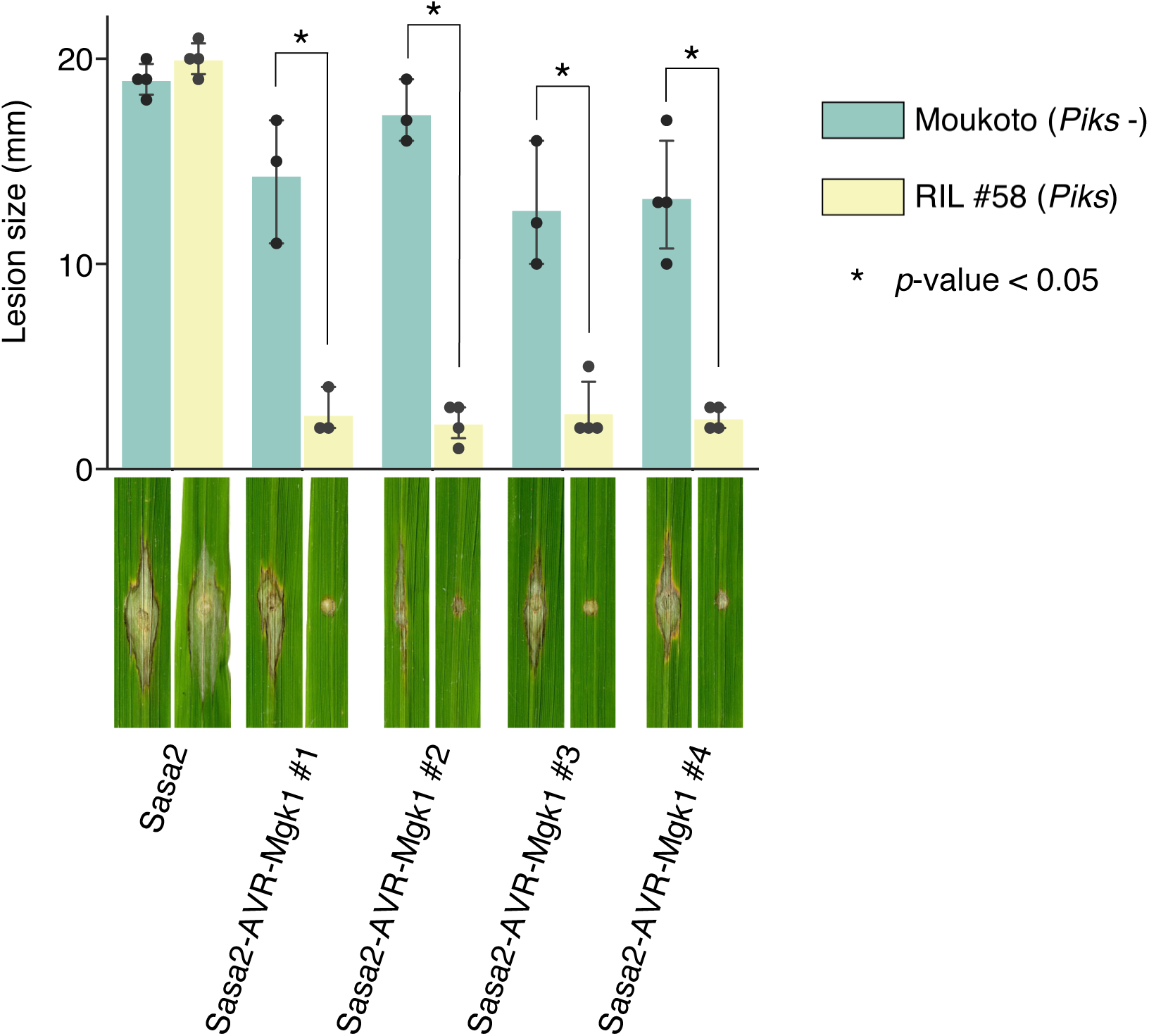
Sasa2 transformants expressing *AVR-Mgk1* cannot infect RIL #58 rice plants containing *Piks*. We produced four independent *M. oryzae* Sasa2 transformants expressing *AVR-Mgk1* and performed punch inoculation assays using wild-type Sasa2 and Sasa2 transformants on rice lines Moukoto (*Piks* -) and RIL #58 (*Piks* +). The lesion size was quantified. Statistically significant differences between rice lines are indicated by asterisks (*p* < 0.05, two-sided Welch’s t-test). The transformant Sasa2-AVR-Mgk1 #4 was used for the punch inoculation assay in **Fig 4D** and **4E** and S9 Fig. The data underlying this figure can be found in **S1 Data**.

**S8 Fig.**
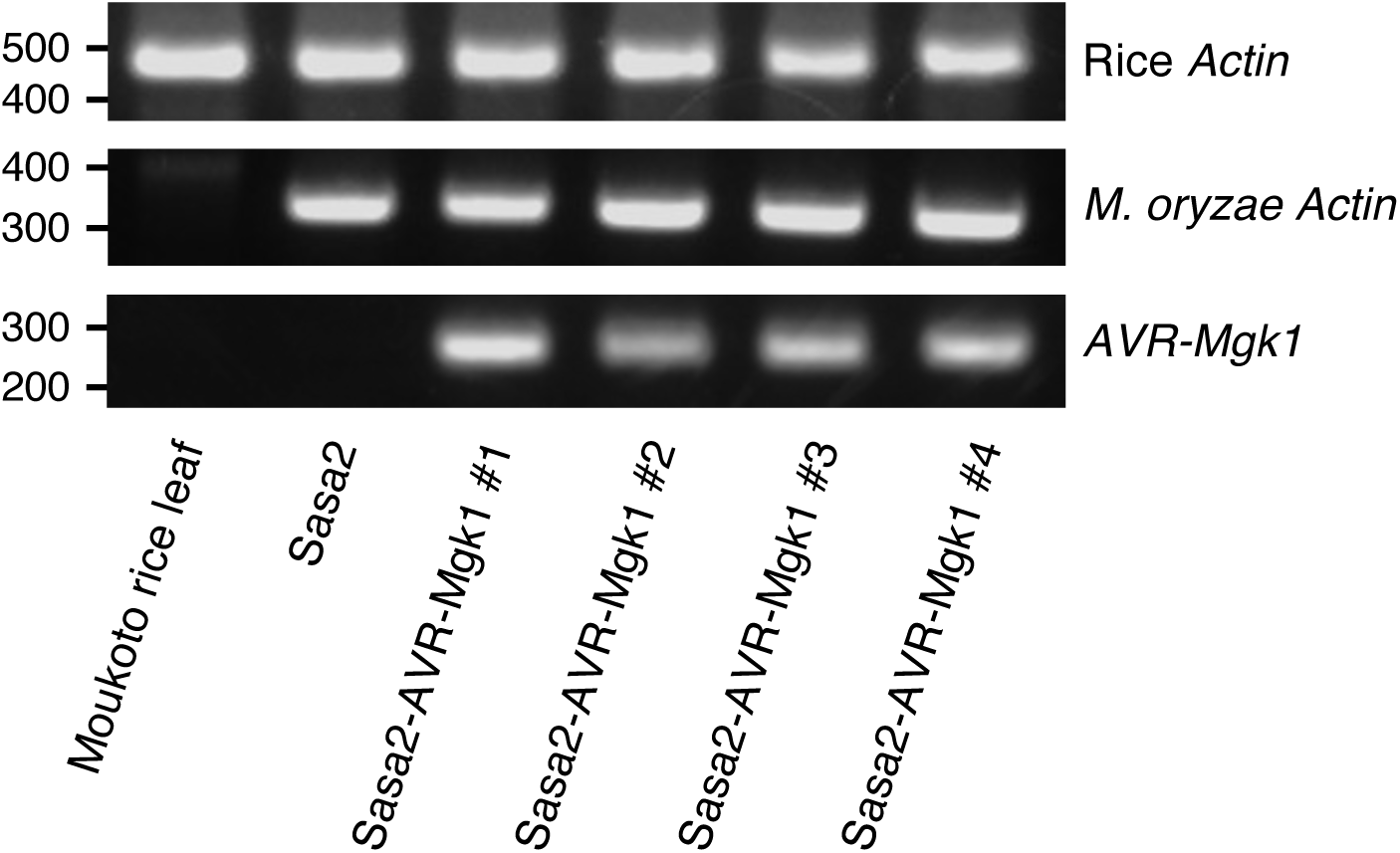
*AVR-Mgk1* expression in infected rice leaves. We punch inoculated independent *M. oryzae* Sasa2 transformants expressing *AVR-Mgk1* on rice cultivar Moukoto. We reverse transcribed cDNA from RNA extracted from the infected rice leaves and amplified *AVR-Mgk1* via PCR. Rice and *M. oryzae Actin* were used as controls.

**S9 Fig.**
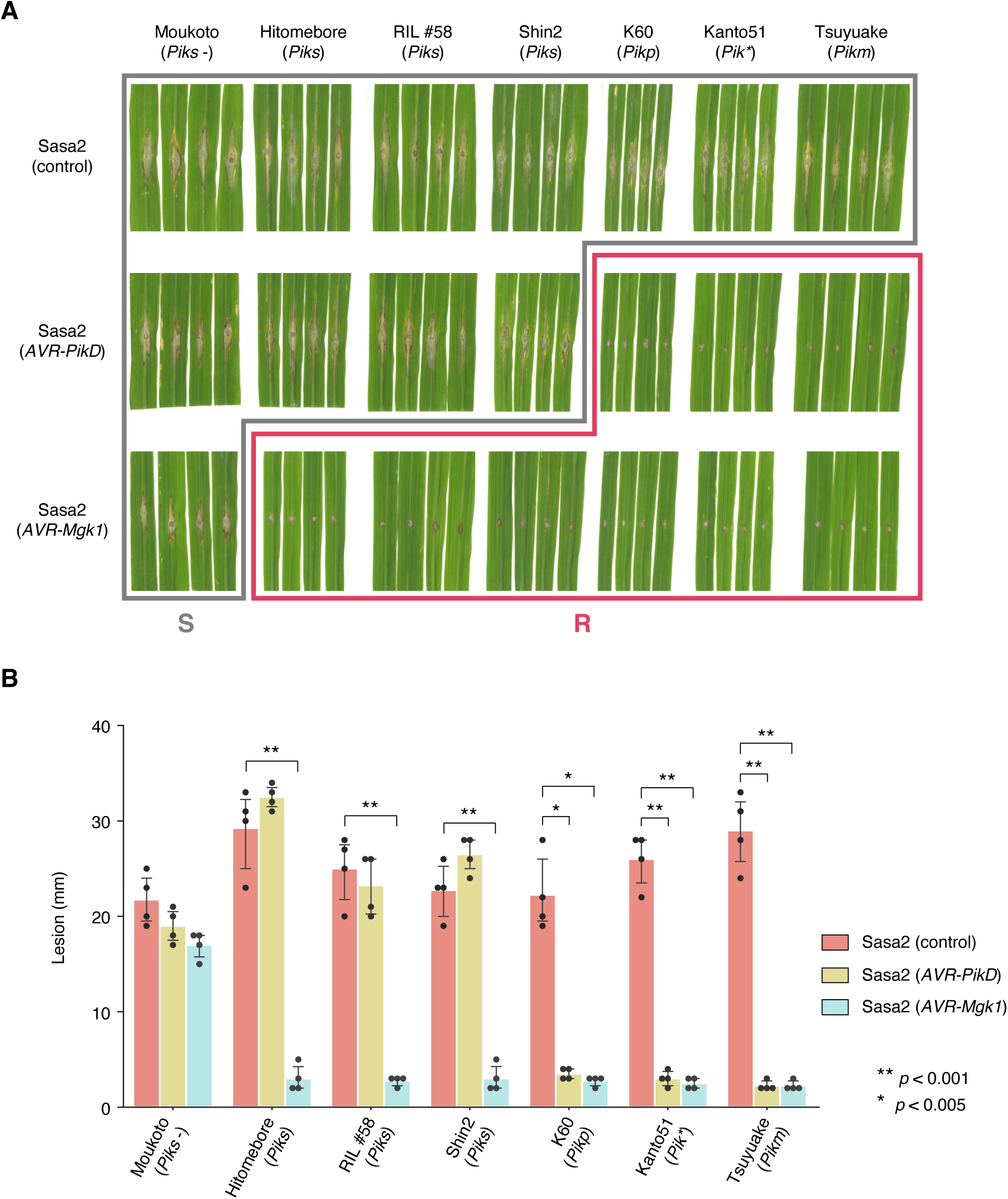
Punch inoculation assays using Sasa2 transformants expressing *AVR-Mgk1* show the broad recognition of AVR-Mgk1 by Pik proteins. (A) We performed punch inoculation assays using wild-type Sasa2 and transformants expressing *AVR-PikD* and *AVR-Mgk1* on rice plants carrying different *Pik* alleles (*Piks*, *Pikp*, *Pik\**, and *Pikm*). A subset of this picture was used in **Fig 4D**. (B) The lesion size in (A) was quantified. Statistically significant differences between isolates are indicated by asterisks (two-sided Welch’s t-test). A subset of this data was used in **Fig 4E**. The data underlying S9B Fig can be found in S1 Data.

**S10 Fig.**
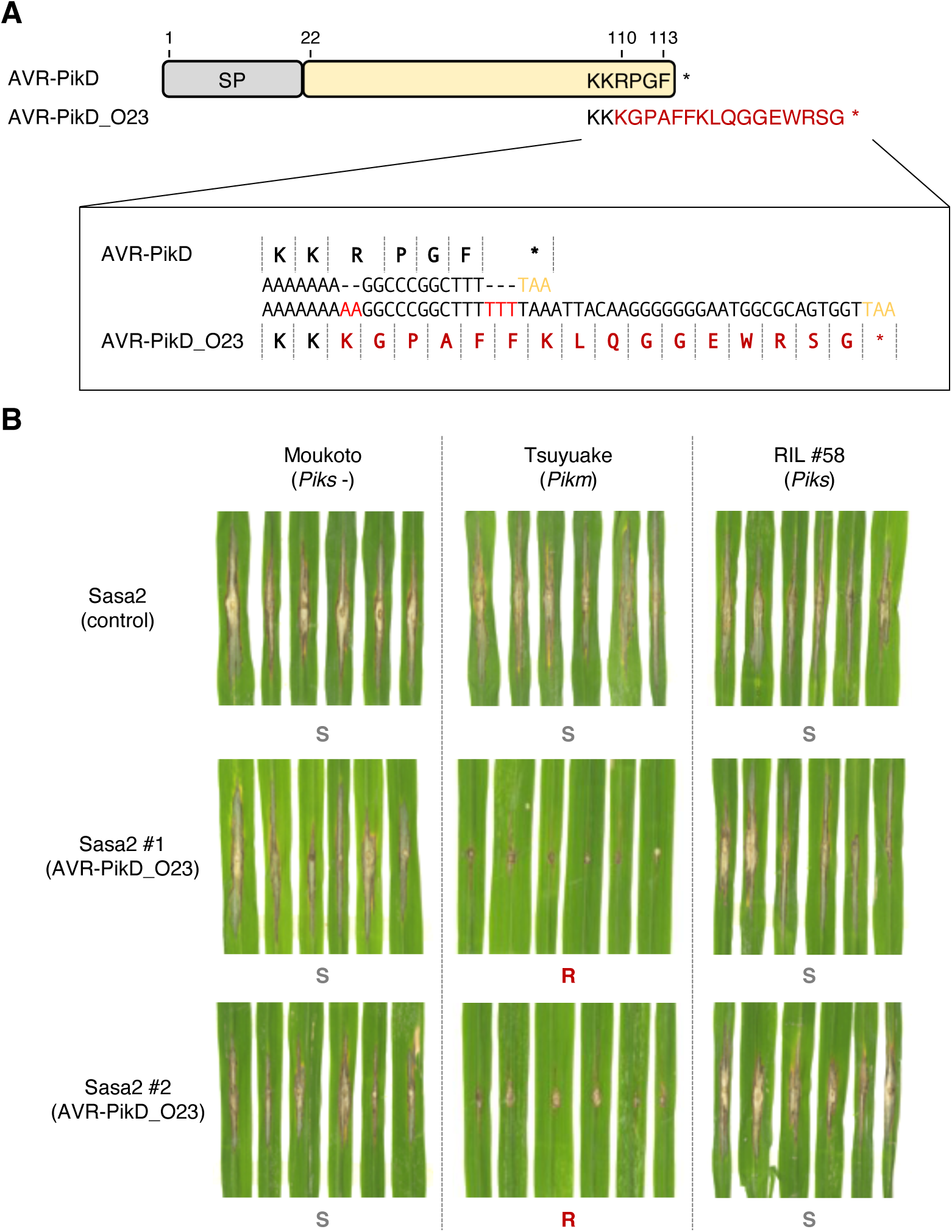
The protein product of *AVR-PikD_O23*, carrying a frameshift mutation, is detected by Pikm but not by Piks. (A) *AVR-PikD* on the O23 mini-chromosome carries a frameshift mutation near the C-terminus that extends the amino acid sequence compared to previously described AVR-PikD. We named this variant *AVR-PikD_O23*. (B) Punch inoculation assays using Sasa2 and its transformant expressing *AVR-PikD_O23*. Sasa2 transformants expressing *AVR-PikD_O23* could not infect Tsuyuake (*Pikm*) but infected RIL #58 (*Piks*).

**S11 Fig.**
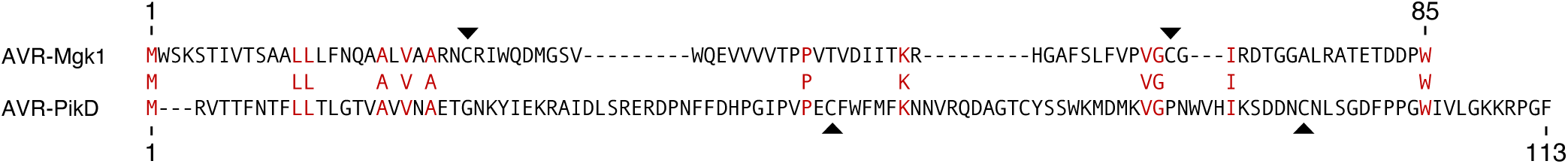
Global sequence alignment reveals that AVR-Mgk1 and AVR-PikD are unrelated in amino acid sequence. We aligned the AVR-Mgk1 and AVR-PikD amino acid sequences using the Needleman-Wunsch global sequence alignment algorithm [158]. Twelve amino acids (red) are identical between AVR-Mgk1 and AVR-PikD. The two cysteine residues conserved in the MAX effector superfamily [28] are indicated by black arrowheads.

**S12 Fig.**
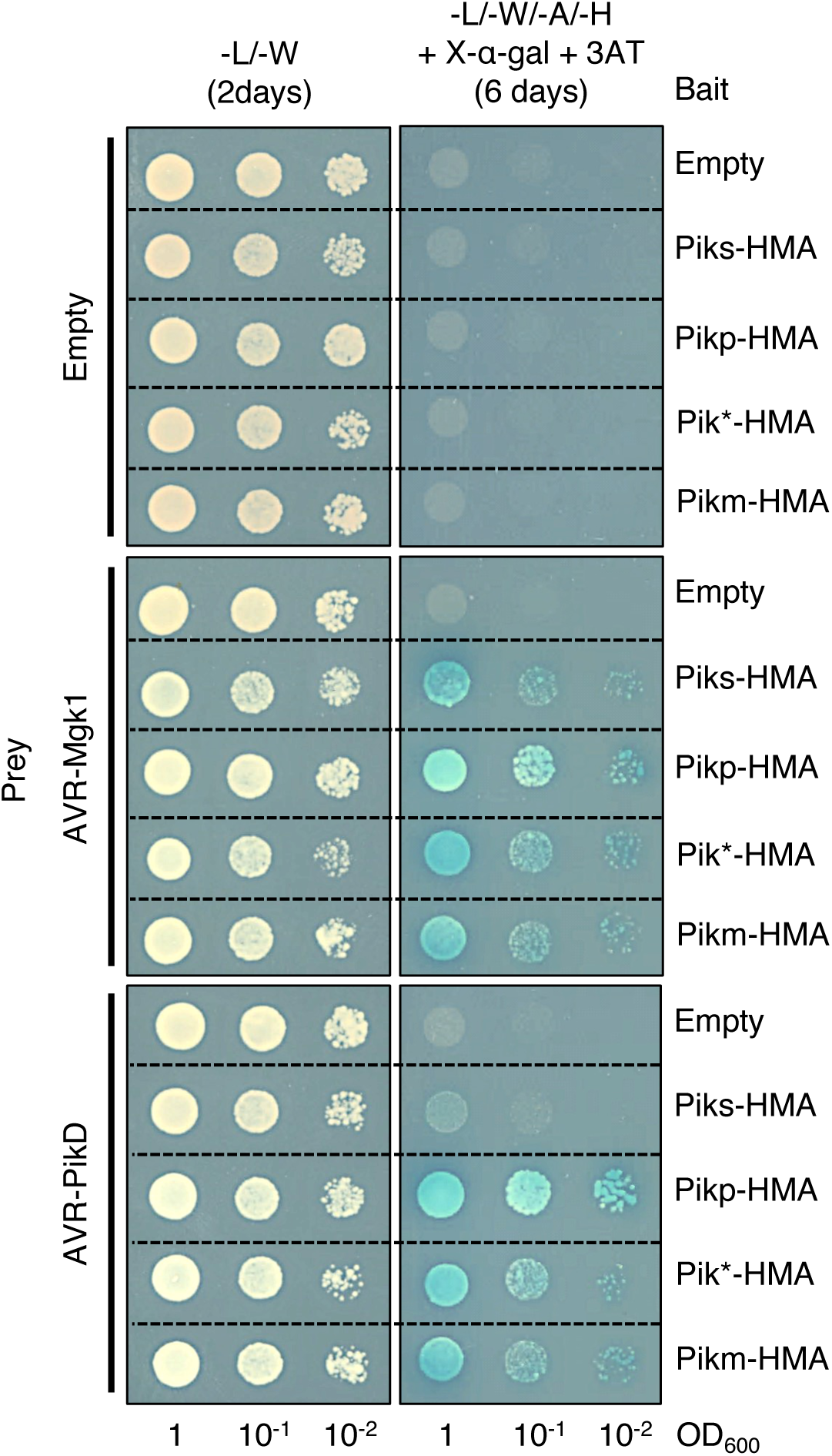
Yeast two-hybrid assay shows that the HMA domains of Pik proteins (bait) bind AVR-Mgk1 (prey). We used HA-tagged AVRs as prey and Myc-tagged HMA domains as bait. Empty vector was used as a negative control. Left side: basal medium lacking leucine (L) and tryptophan (W) for growth control. Right side: basal medium lacking leucine (L), tryptophan (W), adenine (A), and histidine (H) and containing X-α-gal and 10 mM 3AT for selection.

**S13 Fig.**
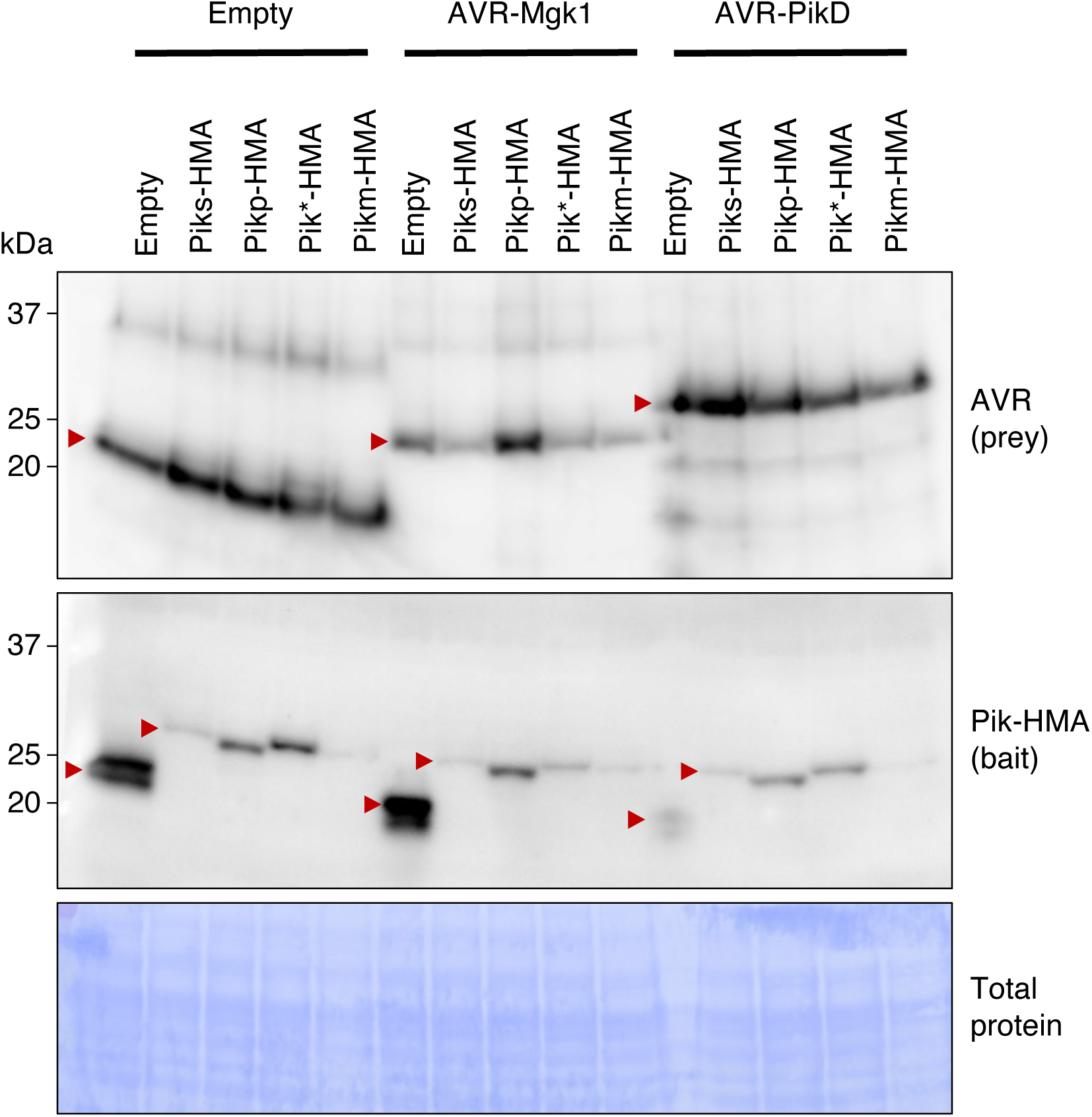
Accumulation of AVRs (prey) and HMA domains (bait) in yeast cells as confirmed by immunoblot analysis. To confirm protein accumulation for the yeast two-hybrid assay, we detected HA-tagged AVRs (prey) by anti-HA antibody and Myc-tagged HMA domains (bait) by anti-Myc antibody. Total proteins of yeast cells detected by Coomassie brilliant blue staining are shown in the bottom as a loading control.

**S14 Fig.**
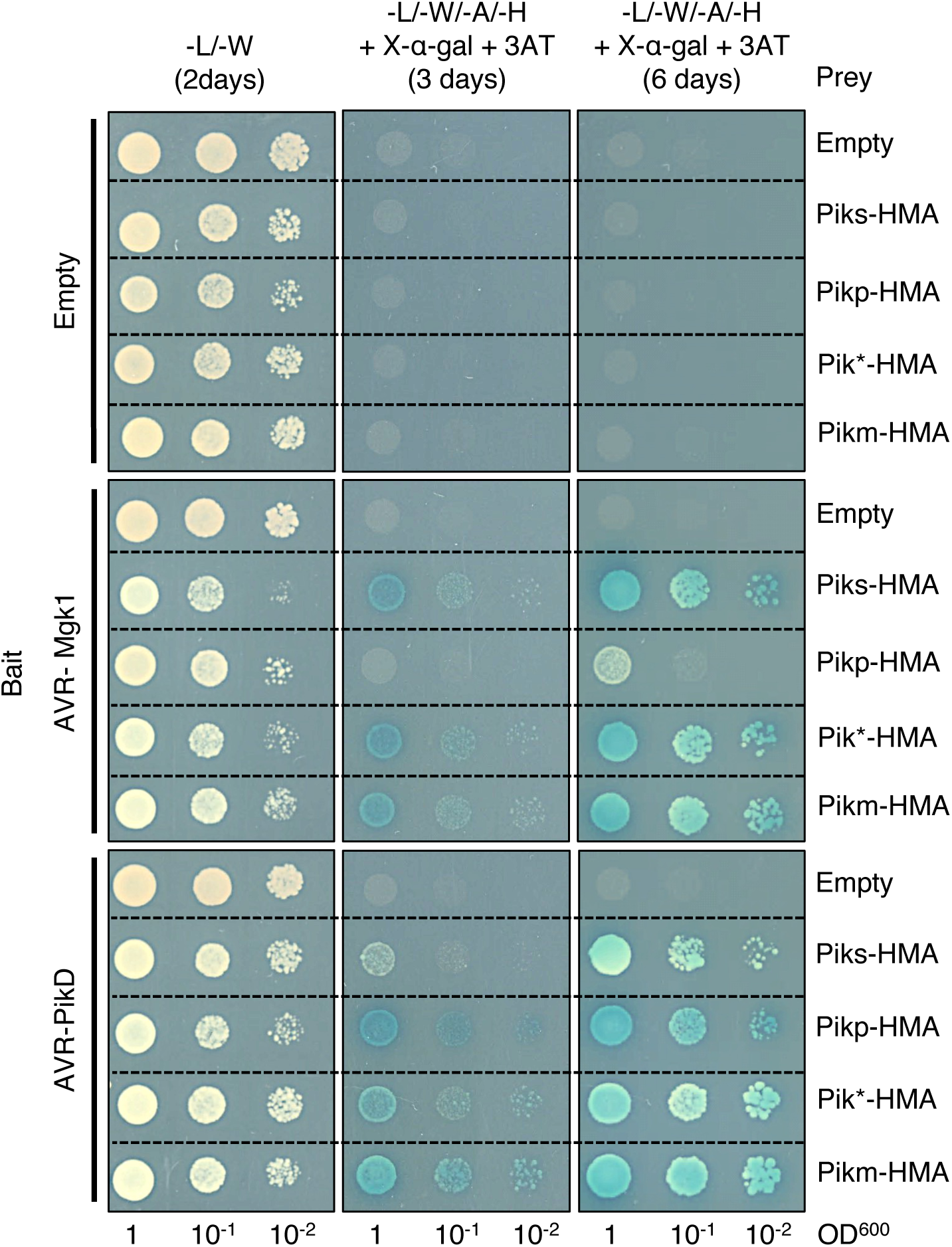
Yeast two-hybrid assay shows that the HMA domains of Pik proteins (prey) bind AVR-Mgk1 (bait). We used Myc-tagged AVRs as bait and HA-tagged HMA domains as prey. Empty vector was used as a negative control. Left side: basal medium lacking leucine (L) and tryptophan (W) for growth control. Right side: basal medium lacking leucine (L), tryptophan (W), adenine (A), and histidine (H) and containing X-α-gal and 10 mM 3AT for selection.

**S15 Fig.**
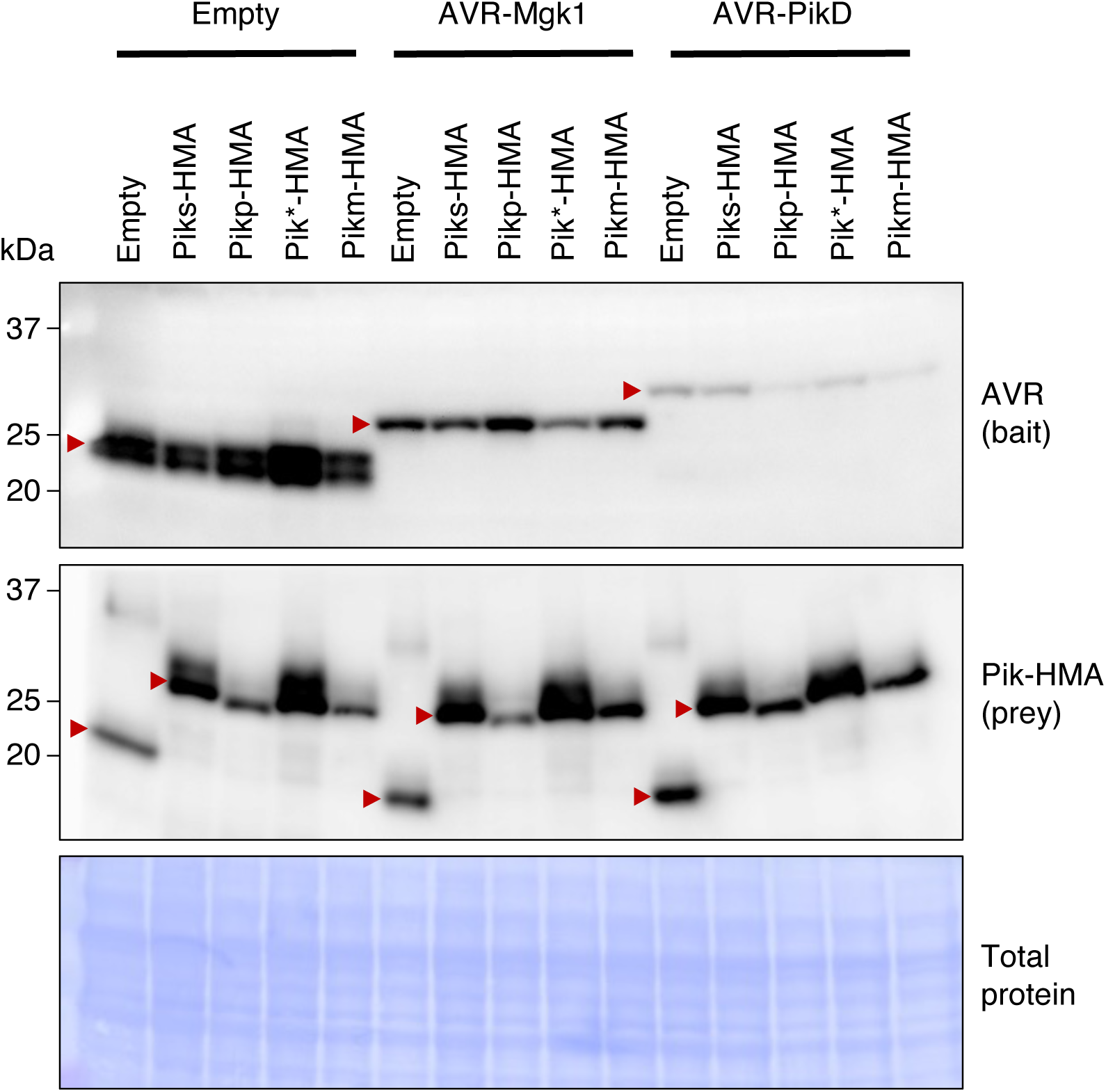
Accumulation of AVRs (bait) and HMA domains (prey) in yeast cells as confirmed by immunoblot analysis. To confirm protein accumulation for the yeast two-hybrid assay, we detected Myc-tagged AVRs (bait) by anti-Myc antibody and HA-tagged HMA domains (prey) by anti-HA antibody. Total proteins of yeast cells detected by Coomassie brilliant blue staining are shown in the bottom as a loading control.

**S16 Fig.**
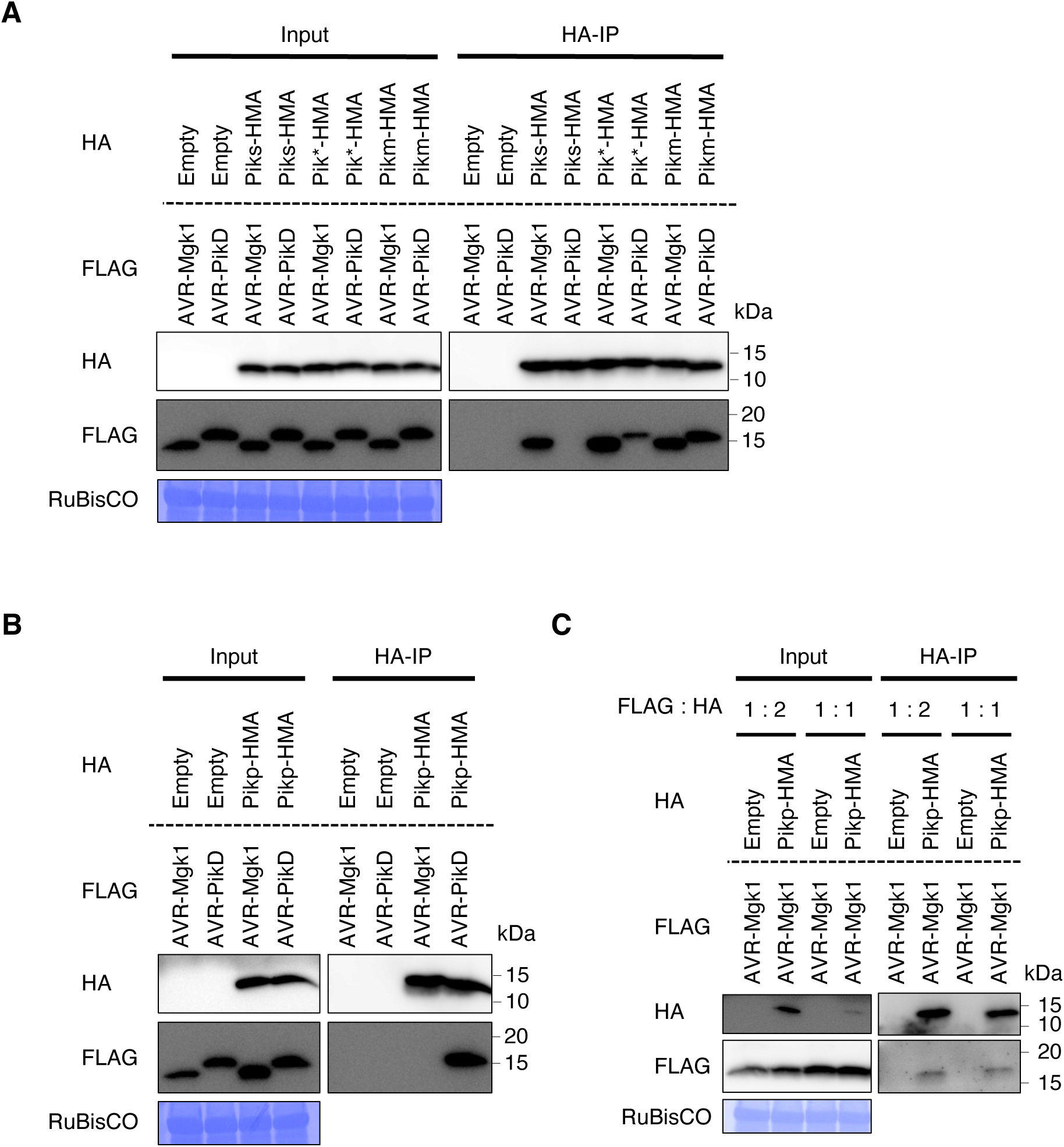
AVR-Mgk1 interacts with the HMA domains of Pik proteins in an *in vitro* co-IP experiment. (A) *In vitro* co-IP experiment between AVR-Mgk1 or AVR-PikD and the HMA domains of Piks (Piks-HMA), Pikm (Pikm-HMA), or Pik* (Pik*-HMA) (1:4 mixed ratio). (B) *In vitro* co-IP experiment between AVR-Mgk1 or AVR-PikD and the HMA domain of Pikp (Pikp-HMA) (1:4 mixed ratio). (C) *In vitro* co-IP experiment between AVR-Mgk1 and Pikp-HMA (1:2 or 1:1 mixed ratios). N-terminally tagged FLAG:AVRs and HA:HMA were expressed in *N. benthamiana*. Empty vector was used as a negative control. We diluted the lysates of AVRs and HMA domains to compare the results at the same concentration and mixed them (1:4, 1:2, or 1:1 ratio) *in vitro* to assemble the protein complex. The protein complexes were pulled down by HA:HMA using Anti-HA affinity gel. *In vitro* co-IP experiments between AVR-Mgk1 and Pikp-HMA (1:2 or 1:1 mixed ratios) were photographed in long-exposure time. The large subunit of ribulose bisphosphate carboxylase (RuBisCO) stained by Coomassie brilliant blue is shown as a loading control.

**S17 Fig.**
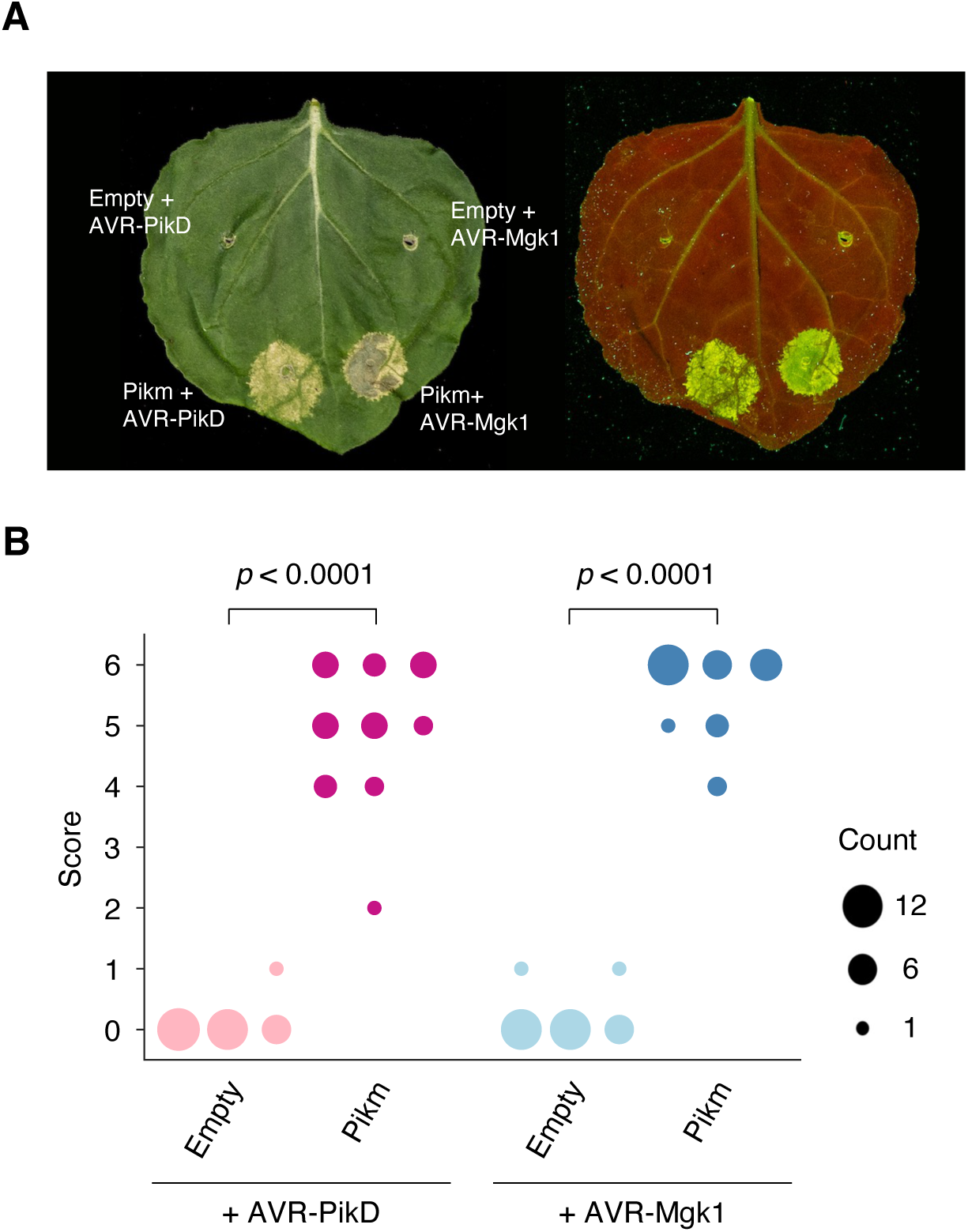
AVR-Mgk1 and AVR-PikD alone do not trigger the HR in *N. benthamiana*. (A) Representative images 5–6 days after transiently co-expressing AVR-Mgk1 and AVR-PikD, either with an empty vector control only expressing p19 or with Pikm, respectively, in *N. benthamiana*. The leaves were photographed under daylight (left) and UV light (right). (B) We quantified the HR in (A) and statistically significant differences are indicated (Mann-Whitney U rank test). Each column shows an independent experiment. The data underlying **S17B Fig** can be found in **S1 Data**.

**S18 Fig.**
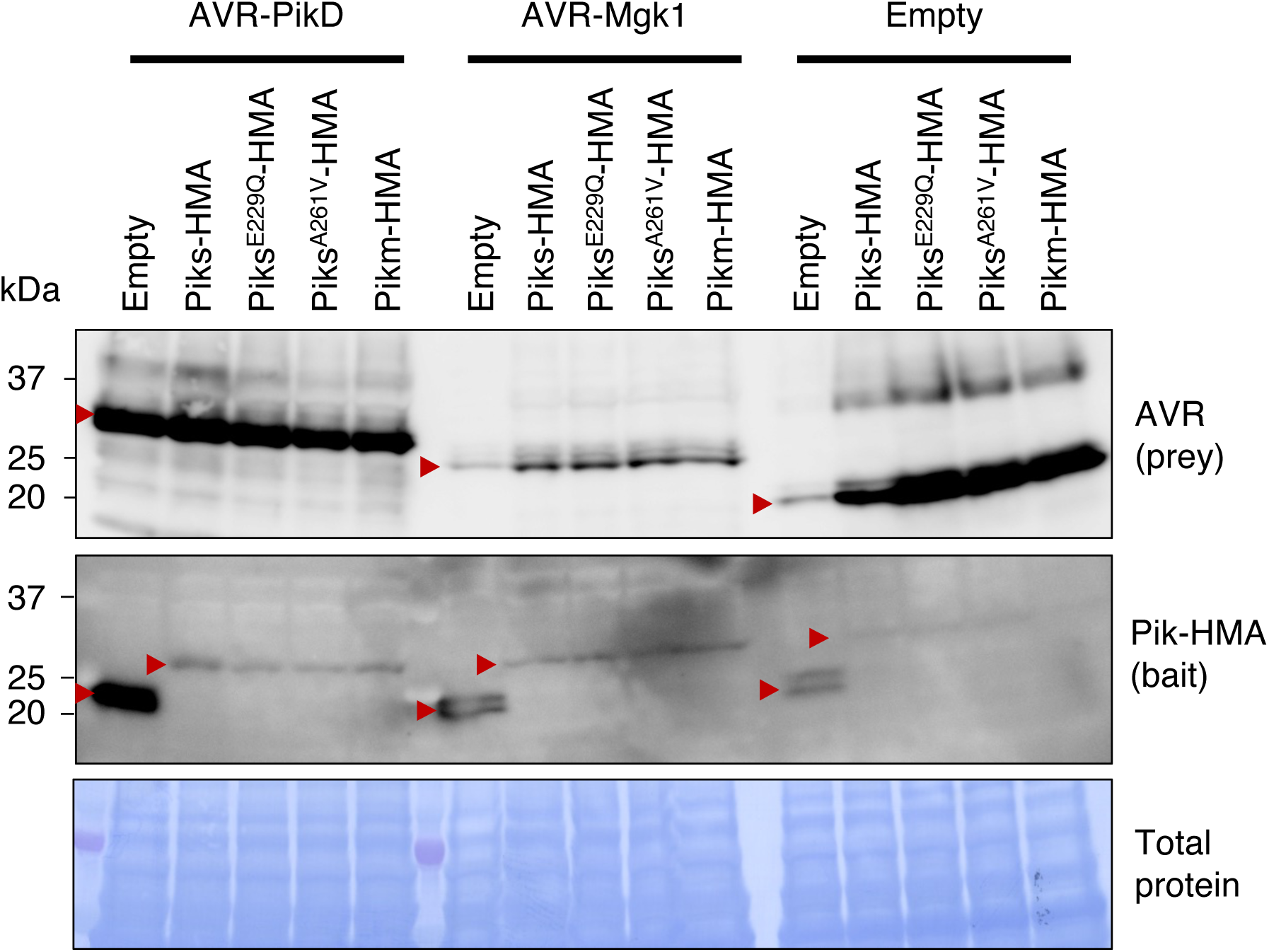
Accumulation of AVRs (prey) and Piks-HMA mutants (bait) in yeast cells as confirmed by immunoblot analysis. To confirm protein accumulation for the yeast two-hybrid assay (**Fig 9A**), we detected HA-tagged AVRs (prey) by anti-HA antibody and Myc-tagged HMA domains (bait) by anti-Myc antibody. Total proteins of yeast cells detected by Coomassie brilliant blue staining are shown in the bottom as a loading control.

**S19 Fig.**
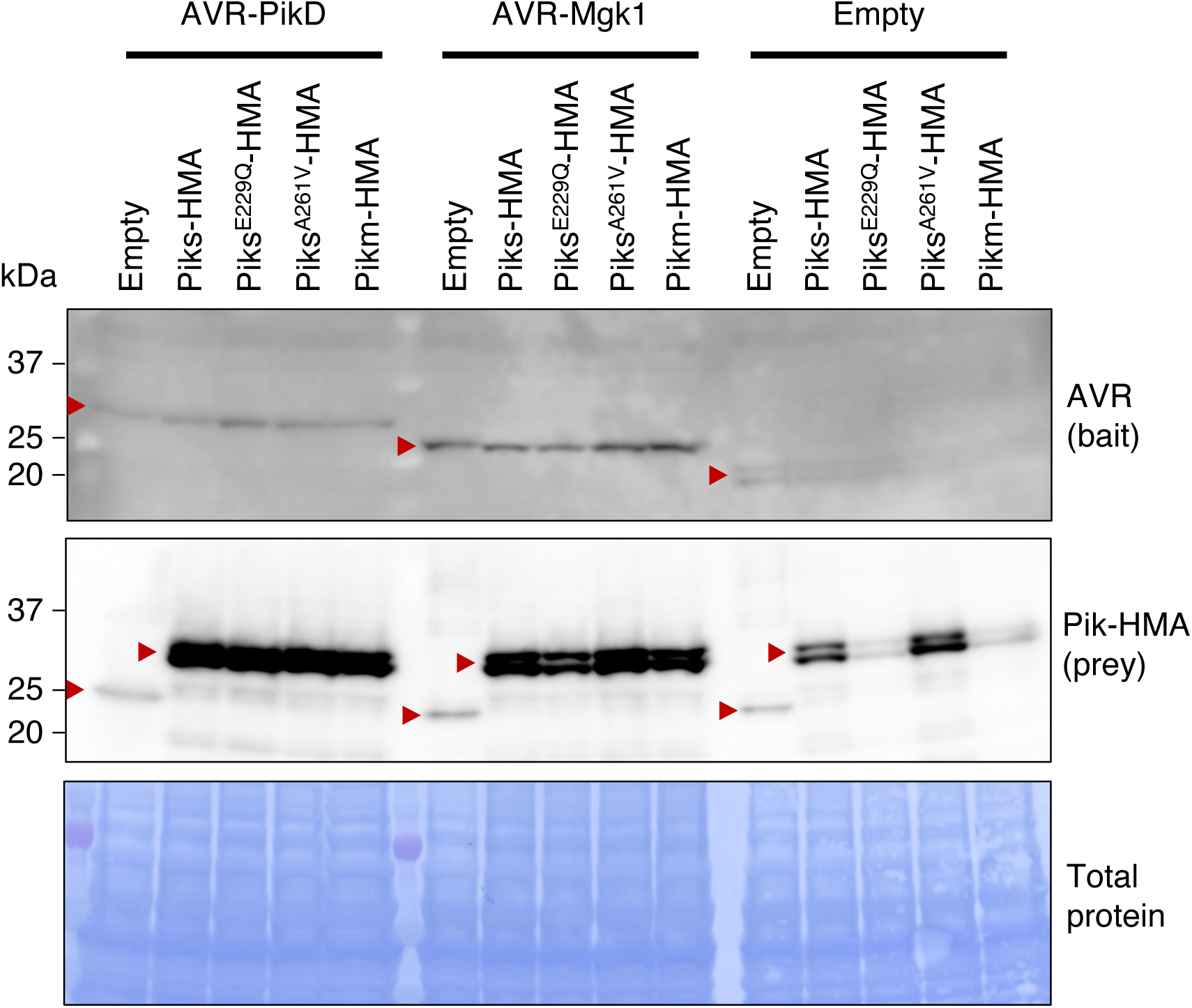
Accumulation of AVRs (bait) and Piks-HMA mutants (prey) in yeast cells as confirmed by immunoblot analysis. To confirm protein accumulation for the yeast two-hybrid assay (**Fig 9B**), we detected Myc-tagged AVRs (bait) by anti-Myc antibody and HA-tagged HMA domains (prey) by anti-HA antibody. Total proteins of yeast cells detected by Coomassie brilliant blue staining are shown in the bottom as a loading control.

**S1 Table. Summary of sequences of rice cultivars Hitomebore and Moukoto and their RILs.**

**S2 Table. Summary of phenotypes of rice RILs derived from a cross between Hitomebore and Moukoto.** The scores 0, 1, and 2 indicate resistant, intermediate, and susceptible phenotypes, respectively. We used these scores as a trait in the genetic association analyses (**Fig 2B** and **C**).

**S3 Table. Summary of *de novo* assemblies of the *M. oryzae* isolates TH3, O23, and d44a.** We sequenced TH3o using PacBio and Illumina DNA sequencers, and O23 and the F_1_ progeny d44a using Oxford Nanopore Technologies (ONT) and Illumina DNA sequencers. The *Sordariomyceta* odb9 dataset was used in BUSCO analysis [99].

**S4 Table. Summary of sequences of *M. oryzae* isolates TH3o and O23 and their F_1_ progeny.**

**S5 Table. Summary of infectivity of the TH3o x O23 F_1_ progeny on RIL #58 and Moukoto rice plants.**

**S6 Table. BLAST search results using AVR-Mgk1 as query.** We used different NCBI databases and algorithms to find sequences related to AVR-Mgk1. We did not find hits in the non-redundant (nr) nucleotide collection using BLASTN search, but found one sequence in the non-redundant (nr) protein collection using BLASTP search. We found other sequences related to AVR-Mgk1 in whole-genome shotgun contigs (wgs) of *Magnaporthe* (taxid: 148303) using BLASTN search. All the BLAST searches were performed using default parameters.

**S7 Table. Summary of the various interactions and phenotypes between Pik NLRs and AVRs in this study.** +++ indicates strong, ++ indicates medium, and + indicates weak interactions or phenotypes, while “-” indicates no interactions or phenotypes for the respective experiments performed.

**S8 Table. Primer sequences used in this study.**

**S9 Table. Cloning details of constructs used for the hypersensitive response cell death assays.**

**S1 Data. Underlying numerical data for Figs 2B, 2C, 2D, 4B, 4C, 4E, 5C, 6B, 6E, 7D, 8B, 8C, and 8D, and S2, S4, S5, S6, S7, S9B, and S17B Figs.**

**S1 Raw images. Raw images for Fig 7B and S8, S13, S15, S16, S18, and S19 Figs.**

